# Structural insights into fosfomycin efflux by a streptococcal ABC transporter

**DOI:** 10.64898/2026.07.07.736899

**Authors:** Atsushi Taguchi, Junso Fujita, Mikio Tanabe, Daisuke Takaya, Kazuo Harada, Toshio Moriya, Kaori Fukuzawa, Keiichi Namba, Kunihiko Nishino

## Abstract

Gram-positive bacteria encode a broad array of ABC transporters that mediate substrate translocation across the cell membrane, with some contributing to their survival under environmental stresses such as antimicrobial exposure. While several of these transporters have been shown to exhibit multidrug efflux activity, the functional roles of many others remain unknown. Here, using an efflux pump screen in the opportunistic human pathogen *Streptococcus pneumoniae*, we identified a previously uncharacterized type IV ABC transporter (FoeAB) that confers resistance to the antibiotic fosfomycin. We show that purified FoeAB mediates fosfomycin transport in a liposome-reconstituted system and provide evidence that it functions as a multidrug efflux pump with substrate preferences distinct from those of known efflux pumps. Furthermore, we present cryogenic electron microscopy (cryo-EM) structures of FoeAB in inward- and outward-facing states, which reveal conformational changes associated with nucleotide binding and identify residues important for substrate transport. Collectively, these findings expand the known repertoire of antibiotic-exporting ABC transporters in Gram-positive bacteria and provide structural insight into its transport mechanism.

**SIGNIFICANCE:** Efflux pumps often play major roles in antibiotic resistance in clinically important pathogens, yet many of these transporters remain poorly defined. In this study, we identify and mechanistically characterize FoeAB, a previously unrecognized type IV ABC transporter in streptococcal species that mediates efflux of the antibiotic fosfomycin. By combining functional assays with structural studies of FoeAB in multiple conformational states, we propose a model for how this transporter recognizes its substrate and harnesses nucleotide-driven conformational changes to mediate substrate export. Our findings point to a previously underappreciated efflux capabilities in Gram-positive bacteria, highlighting the need for better characterization of membrane transporters to clarify their roles in bacterial physiology and drug resistance.

## INTRODUCTION

A key trait that enables bacteria to adapt and proliferate in dynamic environments is their ability to actively export various molecules, including toxic compounds, from the cell. This process is mediated by membrane-bound exporters collectively known as efflux pumps. While these pumps participate in diverse cellular processes, such as biofilm formation and virulence, their role in antibiotic export has received considerable attention due to the growing threat of antimicrobial resistance (1–3). Multidrug resistance phenotypes found in many Gram-negative clinical isolates have been linked to resistance-nodulation-cell division (RND)-type efflux pumps, which extrude a broad range of substrates from both the cytoplasm and periplasm through a tripartite complex spanning the inner and outer membranes (4). In contrast, Gram-positive bacteria lack an outer membrane and do not possess classical RND pumps; however, these bacteria still encode efflux pumps belonging to the major facilitator superfamily (MFS), the multidrug and toxin extrusion (MATE) family, the small multidrug resistance (SMR) family, and the ATP-binding cassette (ABC) family (5).

Among these efflux pumps, several ABC transporters have been linked to antimicrobial resistance in clinically important pathogens (6–8). ABC transporters are primary active transporters sharing a common architecture consisting of two transmembrane domains (TMDs) coupled to two nucleotide-binding domains (NBDs). Unlike their eukaryotic counterparts, which are often encoded as a single polypeptide chain, prokaryotic ABC exporters generally consist of two half-transporters, each containing one TMD and one NBD, that assemble into either a homodimer or a heterodimer (9). Many of the ABC transporters involved in antimicrobial efflux belong to the type IV family based on the recent classification schemes (10). The general conformational cycle of heterodimeric type IV ABC exporters has been established through structural studies of model transporters such as *Thermus thermophilus* TmrAB (11, 12). In the basal, inward-facing state, the TMDs form a cytoplasm-accessible cleft that accommodates the substrate. In TmrAB, ATP binding to both NBDs has been shown to induce NBD dimerization and the transition to an outward-facing conformation, exposing the substrate-binding site to the extracellular side and allowing its transport. ATP hydrolysis and phosphate release promote the return to the inward-facing state, completing the alternating-access cycle.

To date, most characterized Gram-positive multidrug ABC exporters have been found to exhibit broadly overlapping substrate preferences (13, 14). Fluoroquinolone antibiotics are among the shared substrates of these transporters, and their overexpression reduces cellular sensitivity to fluoroquinolones (15). Other substrates range from the anticancer drug daunorubicin to Hoechst dyes, which are also recognized by eukaryotic multidrug ABC transporters, exemplified by the well-characterized human P-glycoprotein (ABCB1) (16, 17). Despite their chemical diversity, these substrates typically contain a planar aromatic motif. The importance of this motif has recently been highlighted by structural studies of the *Bacillus subtilis* multidrug ABC exporter BmrCD in complex with Hoechst dyes, which revealed multiple aromatic interactions between the transporter and substrate that are critical for substrate recognition (18, 19). In contrast, little is known about Gram-positive ABC efflux pumps that transport compounds outside this commonly described set of substrates, including their potential roles in antibiotic efflux and the molecular basis of substrate recognition.

Here, we focused on identifying efflux pumps capable of transporting antimicrobial compounds in the Gram-positive opportunistic pathogen *Streptococcus pneumoniae*. *S. pneumoniae* is responsible for respiratory tract infections that can progress to severe diseases such as pneumonia and meningitis, and it is one of the leading causes of deaths associated with antibiotic resistance (20, 21). The heterodimeric multidrug ABC exporter PatAB has been shown to confer fluoroquinolone resistance in clinical strains, raising the question of whether additional efflux pumps could also contribute to antimicrobial resistance in this pathogen (15). Our pneumococcal efflux pump screen uncovered a previously uncharacterized heterodimeric ABC transporter, FoeAB (<u>Fo</u>sfomycin <u>e</u>fflux pump A and B), that conferred resistance to fosfomycin, an antibiotic that inhibits the first cytoplasmic step of cell wall synthesis (22, 23). Fosfomycin is a phosphonic acid that lacks an aromatic ring, making it chemically distinct from the substrates typically transported by known multidrug efflux pumps. Notably, fosfomycin has not previously been reported as a substrate of an efflux pump, suggesting that FoeAB recognizes substrates through interactions distinct from those of previously characterized multidrug ABC exporters. To elucidate the structural basis of this unique substrate preference, we used cryo-EM to determine inward- and outward-facing structures of a FoeAB homolog from *Streptococcus thermophilus*, which revealed key residues involved in substrate transport. Importantly, we present direct evidence of fosfomycin transport by FoeAB using a proteoliposome reconstitution system.

## RESULTS

### Identification of a streptococcal ABC transporter as a putative fosfomycin exporter

*S. pneumoniae* possesses a diverse set of efflux pumps, but only a few have been functionally characterized (24–26). To better understand their role in drug efflux, we performed a systematic screen of pneumococcal efflux pumps involved in antimicrobial resistance. We reasoned that an efflux pump overexpression library would be particularly suitable, as some efflux pumps may be poorly expressed under standard laboratory conditions (27). We prepared 30 *S. pneumoniae* strains carrying putative efflux pump genes encompassing different families under a constitutive promoter (*SI Appendix*, Table S1). These strains were then tested against a panel of antimicrobial compounds on solid media to identify those with increased resistance (Fig. 1A and *SI Appendix*, Fig. S1). As expected, overexpression of the multidrug efflux pump PatAB allowed better survival in the presence of acriflavine, berberine, ciprofloxacin, norfloxacin, and tetracycline, which all contain an aromatic motif. Another ABC transporter, BceAB, which confers resistance to the polypeptide antibiotic bacitracin via target protection, was also identified as a hit (28). These mutants displayed a 2–4 fold increase in minimum inhibitory concentration (MIC) values for the corresponding compounds in liquid media (*SI Appendix*, Table S2). Together, these results demonstrate that the pneumococcal efflux pump overexpression library is an effective tool for identifying transporters involved in antimicrobial resistance.

**Figure 1:**
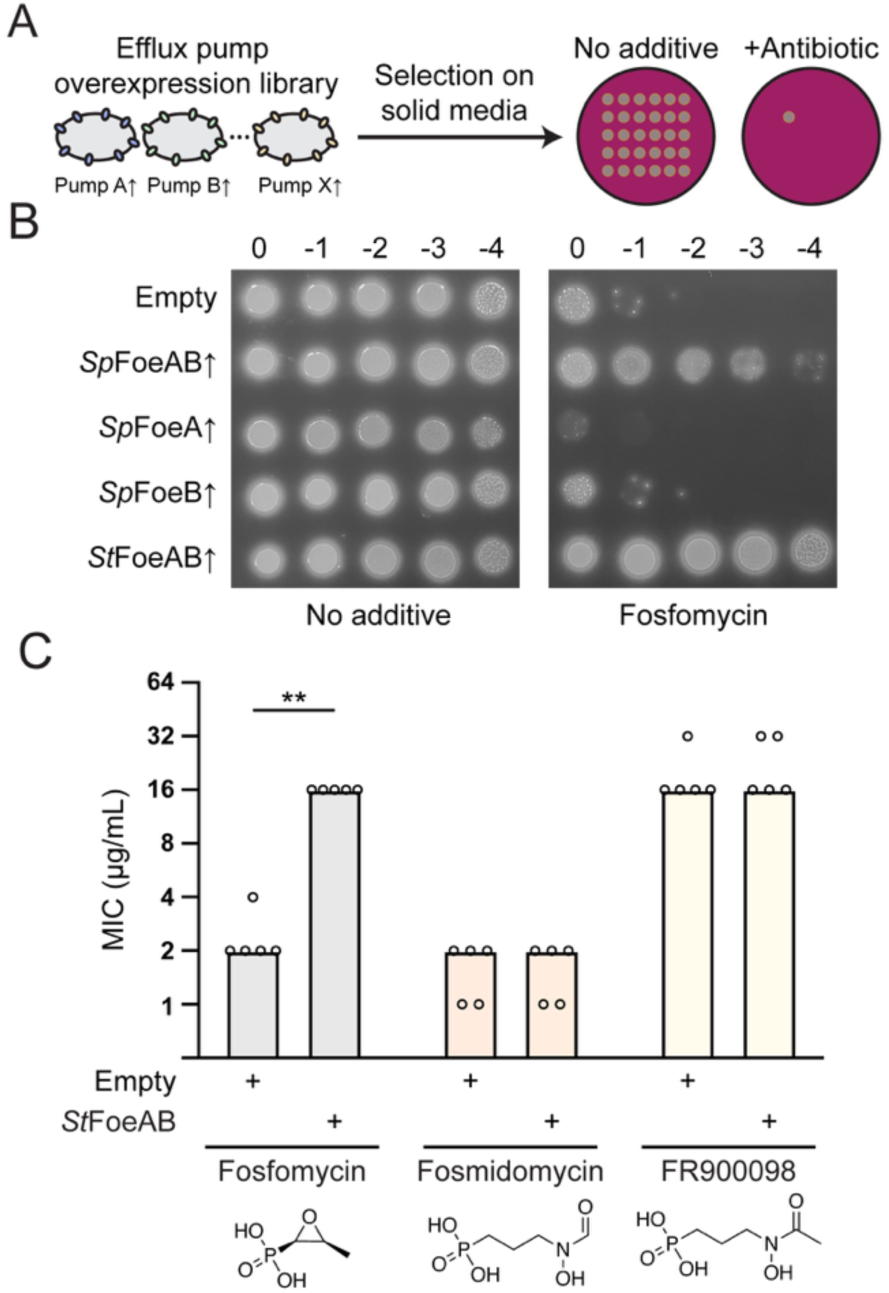
FoeAB overexpression confers fosfomycin resistance. (A) Schematic of the antibiotic efflux pump screen using an *S. pneumoniae* efflux pump overexpression library. Each strain overexpresses a putative efflux pump, and 30 strains were spotted on antibiotic-containing agar plates to assess resistance phenotypes. (B) *S. pneumoniae* strains expressing the indicated gene(s) from a constitutive promoter were spotted on an agar plate containing fosfomycin (50 µg/mL). Strains overexpressing both FoeA and FoeB showed increased resistance against fosfomycin. (C) MIC values for *E. coli* C43(DE3) Δ*acrB* strains harboring either an empty vector or a plasmid expressing FLAG-tagged *St*FoeAB against the indicated phosphonic acid antibiotics. Bars indicate median MIC values, and dots represent individual biological replicates (*n* = 5). \*\**P* < 0.01 (two-sided Mann–Whitney U test).

In addition to the known pneumococcal ABC transporters associated with antimicrobial resistance, we identified an uncharacterized ABC transporter FoeAB (encoded by *spr1216* and *spr1215*) whose overexpression conferred resistance to fosfomycin (Fig. 1B). Fosfomycin targets cell wall synthesis, but unlike many cell wall antibiotics that disrupt later steps of this pathway in the extracellular space, it inhibits the first committed reaction of cell wall precursor synthesis in the cytoplasm, making it a plausible candidate for direct efflux by FoeAB (22, 23). To better characterize FoeAB function, we conducted a series of cell-based assays. Given that the genes encoding FoeAB are in the same operon, we hypothesized that the observed resistance phenotype requires these proteins to form an active heterodimeric complex. We found that *S. pneumoniae* cells overexpressing either FoeA or FoeB alone did not exhibit the same level of fosfomycin resistance as cells overexpressing both proteins, consistent with complex formation (Fig. 1B). FoeAB is broadly conserved among streptococcal species, and we confirmed that this activity is not specific to *S. pneumoniae* FoeAB by showing that overexpression of *Streptococcus thermophilus* FoeAB (*St*FoeAB) also conferred fosfomycin resistance (Fig. 1B).

We next examined whether FoeAB requires additional pneumococcal proteins for its function. If FoeAB activity depended on a native accessory factor, such as a substrate-binding or regulatory protein, we would expect little change in fosfomycin sensitivity when expressed in an unrelated host. We found that *St*FoeAB could be stably expressed in *Escherichia coli* using an isopropyl β-D-thiogalactopyranoside (IPTG)-inducible system, and its expression was sufficient to confer fosfomycin resistance, indicating that other pneumococcal proteins are not essential for FoeAB activity (Fig. 1C). We then used this system to test whether *St*FoeAB protects against other phosphonic acid antibiotics that are active against *E. coli*. Fosmidomycin and its derivative FR900098 are two such compounds that exert lethality by inhibiting the methylerythritol phosphate pathway for isoprenoid biosynthesis, which is essential in *E. coli* but not in *S. pneumoniae* (29). Overexpression of *St*FoeAB did not protect *E. coli* against either antibiotic, suggesting that FoeAB displays some selectivity in recognizing phosphonic acid compounds (Fig. 1C).

### Structural comparison of inward-facing FoeAB in different nucleotide-binding states

Fosfomycin has not been reported as a substrate of any ABC transporter described to date, which prompted us to investigate the molecular features that enable FoeAB-mediated transport. Using the *E. coli* expression system, we successfully purified *St*FoeAB to homogeneity with detergent *n-*dodecyl-β-D-maltopyranoside (DDM), and the DDM-solubilized samples were subjected to cryo-EM analysis to gain structural insights (*SI Appendix*, Fig. S2-S4). We determined four different inward-facing states from the wild-type *St*FoeAB samples (Fig. 2A and *SI Appendix*, Table S3). *St*FoeAB exists as a heterodimer, with each protomer containing one NBD and one TMD comprising six TM helices, two of which domain-swap into the opposing TMD (*SI Appendix*, Fig. S5). This structural organization is prototypical of type IV ABC transporters (10). Although a subset of type IV ABC transporters has been found to contain an additional domain, such as the extracellular domain in *B. subtilis* BmrCD, no comparable domain is present in FoeAB (18). We obtained three nucleotide-free structures from the additive-free dataset, which showed only minimal overall structural differences (*SI Appendix*, Fig. S6). Unexpectedly, we resolved another structure from the same dataset that included an ADP and Mg^2+^ in FoeA NBD (ADP-bound state), which presumably remained bound throughout the purification process (*SI Appendix*, Fig. S7). Addition of the non-hydrolysable ATP analog adenylyl imidodiphosphate (AMPPNP) and ATP to wild-type *St*FoeAB yielded structures with AMPPNP and ATP bound to FoeA NBD, respectively (Fig. 2A and *SI Appendix*, Fig. S7). The latter structure also contained an ADP in FoeB NBD, which is likely a product of ATP hydrolysis that occurred during sample preparation (*SI Appendix*, Fig. S7).

**Figure 2:**
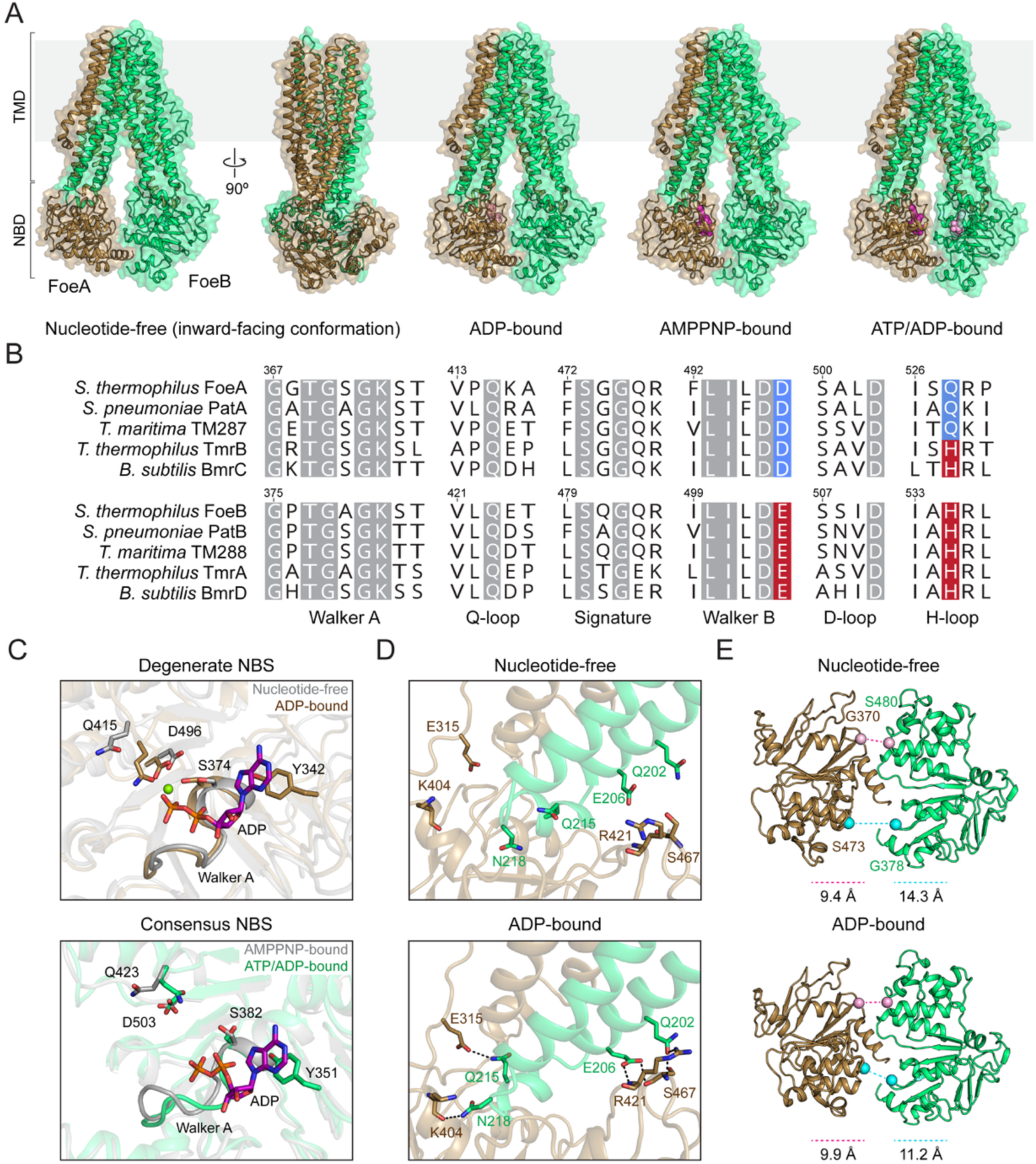
FoeAB is a heterodimeric ABC transporter with asymmetric nucleotide binding sites. (A) Cartoon representations of inward-facing FoeAB in different nucleotide binding states viewed along the membrane plane. Pink and purple spheres correspond to ADP and ATP/AMPPNP, respectively. (B) Sequence alignment of select NBD motifs in FoeAB and representative prokaryotic ABC transporters. Residues conserved in all sequences are shown in gray. The Walker B glutamate and H-loop histidine conserved in the consensus NBS are highlighted in red, while the corresponding aspartate and glutamine replacements observed in the degenerate NBS are highlighted in blue. (C) Close-up views of degenerate and consensus NBSs. The light green sphere indicates the magnesium ion. The Walker A serine (S382^A^), Q-loop glutamine (Q415^A^), and Walker B aspartate (D496^A^) coordinate the magnesium ion in the degenerate NBS. (D) Structural comparison of the FoeA TMD-NBD interface in nucleotide-free and ADP-bound structures. Residues involved in interprotomer interactions that form upon ADP binding are highlighted. (E) Bottom view of the NBDs in nucleotide-free and ADP-bound states. Pink and cyan spheres indicate the signature motif serine and Walker A glycine in the degenerate and consensus NBSs, respectively.

The asymmetrical binding of AMPPNP and ATP to FoeA NBD resembles reported structures of model heterodimeric ABC exporters TmrAB and *Thermotoga maritima* TM287/288 (30, 31). These transporters harbor a degenerate nucleotide-binding site (NBS) that is catalytically impaired due to substitutions in canonical motifs involved in nucleotide binding and/or hydrolysis. To assess whether *St*FoeAB shares this feature, we compared the NBD sequences of *St*FoeAB with these exporters and found that FoeA contains an aspartate substitution of the Walker B catalytic glutamate, a substitution frequently associated with degenerate NBSs. Additionally, the conserved H-loop histidine is replaced with a glutamine in FoeA, which is also present in a subset of degenerate NBSs, suggesting that *St*FoeAB undergoes a catalytic cycle in which only a single NBS is capable of efficient nucleotide hydrolysis (Fig. 2B and *SI Appendix*, Fig. S5).

The asymmetric nature of *St*FoeAB NBSs was evident in the structural reorganization triggered by ADP binding. In both NBSs, ADP is coordinated by Walker A residues that comprise a phosphate-binding loop and a conserved tyrosine that forms a stacking interaction with the adenine ring (Fig. 2C). In the canonical NBS, Mg^2+^ is not present, and FoeB NBD displays only subtle alterations in response to ADP binding. In contrast, in the degenerate NBS, the Q-loop glutamine (Q415^A^) shifts by ∼5 Å, and the Walker A serine (S374^A^) and Walker B aspartate (D496^A^) each shift by ∼2 Å to coordinate Mg^2+^ upon ADP binding. This rearrangement is accompanied by the rotation of the ɑ-helical domain and alteration of the interface between FoeA NBD and FoeB TM helices, where polar residues on TM4 and the subsequent coupling helix stabilize FoeA NBD by forming new electrostatic interactions (Fig. 2D). The conformational changes observed in FoeA NBD reflect a shared asymmetric nucleotide-dependent transition documented in other heterodimeric ABC exporters, and the presence of ADP in the degenerate NBS indicates that γ-phosphate binding is not required for this structural reorganization in *St*FoeAB (12, 31).

One feature that varies among the heterodimeric ABC transporters is the mode of inter-NBD interaction. In *St*FoeAB, nucleotide binding to the degenerate NBS brings the FoeA signature motif closer to the consensus NBS, positioning it for subsequent NBD dimerization (Fig. 2E). This event was proposed to mediate allosteric crosstalk in TM287/288, where new contacts between the D-loop and Walker A motifs are established upon AMPPNP binding; however, no analogous bond formation was observed in *St*FoeAB (*SI Appendix*, Fig. S8) (31, 32). Furthermore, while TmrAB NBDs have been found to form a stable interaction through their C-terminal helices that is critical for its activity, this interaction is absent in *St*FoeAB, highlighting the divergence in inter-NBD communication among ABC exporters (33). Because the observed NBD arrangement could potentially be influenced by detergent solubilization, we further evaluated this possibility by determining an AMPPNP-bound *St*FoeAB structure prepared in peptidiscs, which are amphipathic bi-helical peptides that stabilize membrane proteins in a detergent-free environment (*SI Appendix*, Fig. S9) (34, 35). This structure was near-identical to the AMPPNP-bound structure in DDM, suggesting that NBD separation is not substantially influenced by the surrounding membrane-mimetic environment (*SI Appendix*, Fig. S10).

### The FoeAB inner cavity contains a putative substrate recognition site

The *St*FoeAB TMDs form an inverted V-shaped cavity exposed to the cytoplasm with an estimated volume of ∼4000 Å^3^, sufficient to accommodate molecules of various sizes. Within this cavity, we focused on basic residues that could mediate recognition of the negatively charged fosfomycin phosphonate group and identified a positively charged pocket consisting of K22^A^, R132^A^, and R253^B^ (Fig. 3A). This pocket is readily accessible from the central cavity, and closer inspection of the cryo-EM maps revealed extra density in this region, consistent with a small anion such as phosphate or sulfate (Fig. 3B). Notably, this density was observed under all conditions used to obtain inward-facing structures, regardless of fosfomycin addition, suggesting that the putative ligand represents a co-purified molecule of *E. coli* origin (*SI Appendix*, Fig. S11). Despite extensive attempts to determine a fosfomycin-bound StFoeAB structure, we did not observe clear density attributable to fosfomycin, possibly due to competition between the putative anion and fosfomycin at the pocket. Supporting this idea, molecular docking predicted that the phosphonate group of fosfomycin overlaps with the extra density, with its phosphonate oxygens interacting with the basic pocket residues and its epoxy oxygen contacting R132^A^ (Fig. 3B). To examine the importance of these residues in substrate transport, we evaluated fosfomycin sensitivity of *E. coli* cells expressing *St*FoeAB variants with alanine substitutions in each pocket-forming residue. These alanine variants expressed similarly to wild-type *St*FoeAB but were unable to confer fosfomycin resistance, underscoring the critical role of these residues in fosfomycin transport (Fig. 3C, *SI Appendix*, Fig. S12).

**Figure 3:**
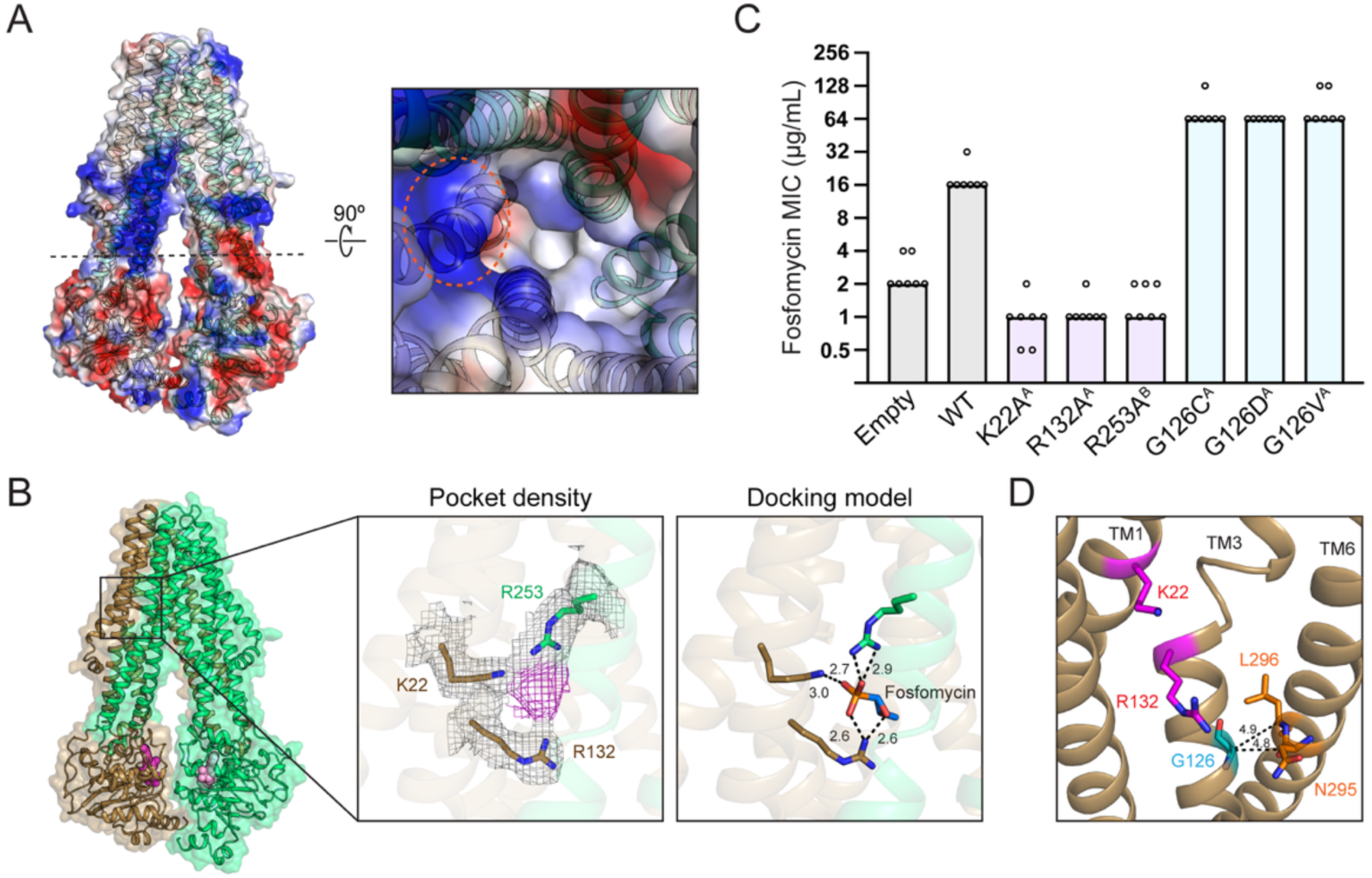
Identification of a putative substrate recognition site in FoeAB. (A) Surface representation of nucleotide-free FoeAB colored according to electrostatic potential. The side view and the interior cavity viewed from the dotted plane are shown. The positively charged pocket is indicated by an orange circle. (B) Close-up view of the positively charged pocket in the ATP/ADP-bound structure. Density corresponding to the basic residues comprising the pocket (gray) and a putative ligand (purple) is contoured at 4σ. A fosfomycin docking model in the pocket is shown in the right panel, with numbers indicating predicted distances between fosfomycin and the pocket residues in angstroms. (C) Fosfomycin MIC values for *E. coli* C43(DE3) Δ*acrB* strains expressing FLAG-tagged FoeAB containing the indicated substitutions. Bars indicate median MIC values, and dots represent individual biological replicates (*n* = 7). Statistically significant differences were observed between wild-type and all variants (*P* < 0.01, Mann–Whitney U tests with Holm-Bonferroni correction). (D) G126^A^ (cyan) is located on TM3 and faces N295^A^ and L296^A^ (orange) on TM6, forming part of the TM3-TM6 interface. K22^A^ and R132^A^ are colored in magenta for clarity.

We next asked whether *St*FoeAB variants with enhanced fosfomycin transport activity could further shed light on the molecular determinants of fosfomycin transport. To identify such variants, we used an untargeted approach in which we generated a *St*FoeAB expression plasmid library carrying random point mutations in TMD residues spanning TM3-6 of FoeA and FoeB. This library was introduced in *E. coli*, and transformants that grew on solid media containing lethal concentrations of fosfomycin were selected for further analysis. This screen yielded *St*FoeAB mutants with substitutions in G126^A^ (G126C, G126D, and G126V), which conferred a 4-fold increase in fosfomycin MIC (Fig. 3C). G126^A^ lies at the TM3-TM6 interface and faces N295^A^ and L296^A^ on the opposing helix, with Cα–Cα distances of approximately 5 Å (Fig. 3D). The recovery of multiple substitutions at this position suggests that the gain-of-function phenotype results from disruption of the native glycine-mediated packing interaction rather than from a specific side-chain chemistry, potentially increasing the conformational flexibility of TM3 and affecting the positioning of R132^A^ for better substrate recognition.

### The outward-facing FoeAB structure provides insight into substrate release

To gain further insight into the *St*FoeAB transport cycle, we next aimed to obtain an outward-facing structure. Previous studies of heterodimeric ABC exporters have used ATPase-impaired mutants in which the consensus Walker B glutamate is replaced with glutamine to stabilize outward-facing states (12, 36, 37). To this end, we introduced an analogous substitution E504Q^B^ in *St*FoeAB and used this mutant to trap the transporter in an ATP/ATP-bound state for cryo-EM analysis (*SI Appendix*, Fig. S13). In line with prior reports, the E504Q^B^ mutant adopted an outward-facing conformation with ATP bound at both NBSs (Fig. 4A). The NBDs form a tight dimer, with the signature motif serine interacting with the ATP γ-phosphate at both NBSs (*SI Appendix*, Fig. S14). In contrast to the ATP/ADP-bound structure, Mg^2+^ was present in the consensus NBS, suggesting that the terminal phosphate plays a pivotal role in Mg^2+^ coordination. NBD dimerization induces a major structural reorganization of the TMD region, which involves the opening of the extracellular gate and the formation of the V-shaped cavity that extends into the cytoplasmic side of the membrane. Newly established contacts between TM3 and TM4 are present in both protomers at the base of this cavity, stabilizing the closure of the intracellular gate (*SI Appendix*, Fig. S15).

**Figure 4:**
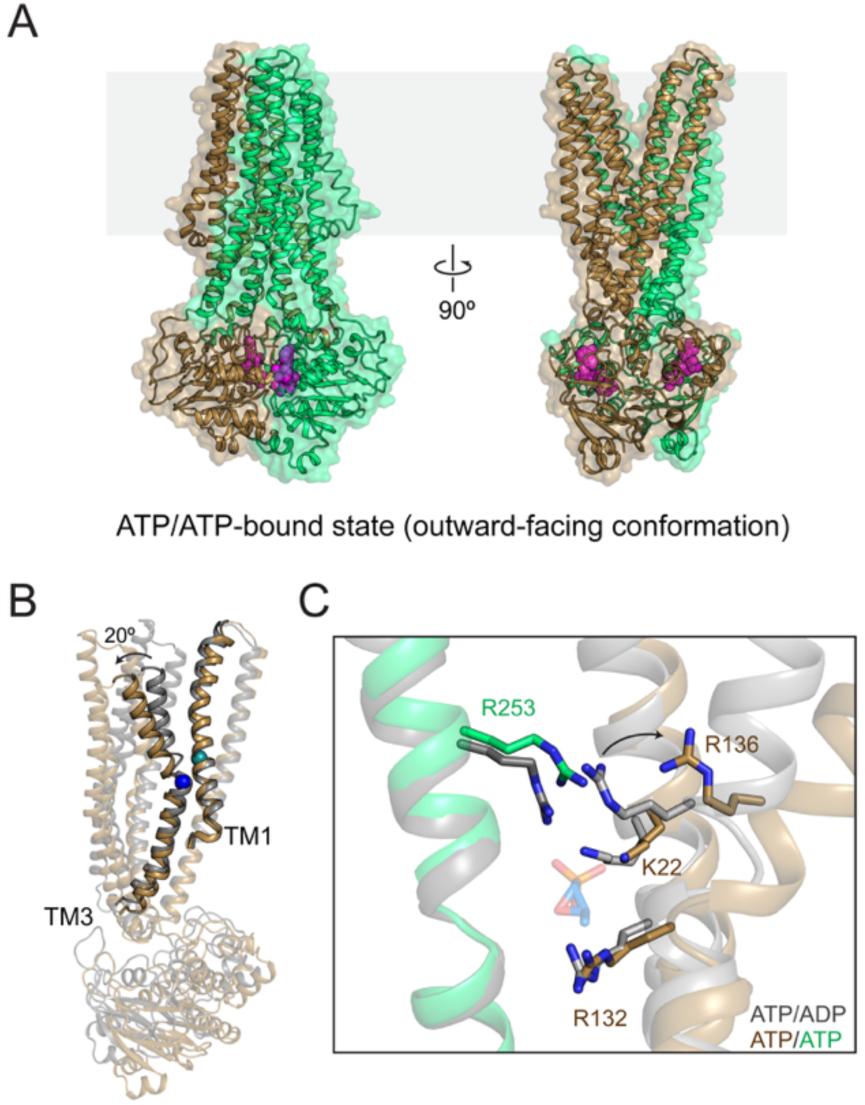
Outward-facing structure of FoeAB suggests a potential mechanism for substrate release. (A) Cartoon representations of outward-facing FoeAB(E504Q^B^) viewed along the membrane plane. Purple spheres represent ATP. (B) Overlay of FoeA in the ATP/ADP-bound (gray) and ATP/ATP-bound (brown) states. Teal and blue spheres correspond to K22^A^ and R132^A^, respectively. (C) Overlay of the putative substrate recognition site in the ATP/ADP-bound (gray) and ATP/ATP-bound (brown/green) states. Fosfomycin from the docking model (blue) is shown for reference. The arrow indicates the movement of R136^A^ between the inward-facing and outward-facing conformations.

The positively charged pocket is exposed to the extracellular side in the outward-facing state, consistent with its proposed role in substrate coordination. Structural comparison of the inward and outward-facing structures revealed only modest changes in the relative C⍺ positions of pocket-forming residues, with R132^A^ moving ∼1 Å away from K22^A^ and R253^B^ upon transitioning to the outward-facing conformation. However, this transition has a profound effect on residues located in the extracellular half of FoeA TM3, which bends 20° outward from R132^A^ (Fig. 4B). One such residue is R136^A^, which forms a parallel stacking arrangement with R253^B^ and sterically restricts its side chain movement in the inward-facing state (Fig. 4C). Separation of these residues in the outward-facing conformation destabilizes this arrangement and allows greater conformational flexibility of R253^B^, likely facilitating substrate release from this pocket by weakening substrate coordination. In agreement with this model, the cryo-EM map of the outward-facing structure lacked the extra density observed in the inward-facing structures (*SI Appendix*, Fig. S16).

### *St*FoeAB transports fosfomycin *in vitro*

Encouraged by the cellular and structural results supporting the role of FoeAB as a fosfomycin exporter, we biochemically analyzed the transport activity of liposome-reconstituted *St*FoeAB. We first assessed the functionality of the wild-type transporter by measuring its ATPase activity and found that *St*FoeAB exhibited high basal activity comparable to that of previously reported prokaryotic multidrug-efflux ABC exporters (Fig. 5A and *SI Appendix*, Table S4) (38). We also examined whether FoeAB could efficiently hydrolyze GTP because the pneumococcal ABC transporter PatAB was reported to favor GTP over ATP as its nucleotide substrate (39). While *St*FoeAB exhibited moderately lower catalytic efficiency for GTP than for ATP, it showed a higher GTP hydrolysis rate under saturating conditions (Fig. 5B and *SI Appendix*, Table S4). These results suggest that FoeAB may use GTP as an alternative nucleotide substrate, especially in *S. pneumoniae* cells where the abundance of ATP and GTP was found to be similar (39).

**Figure 5:**
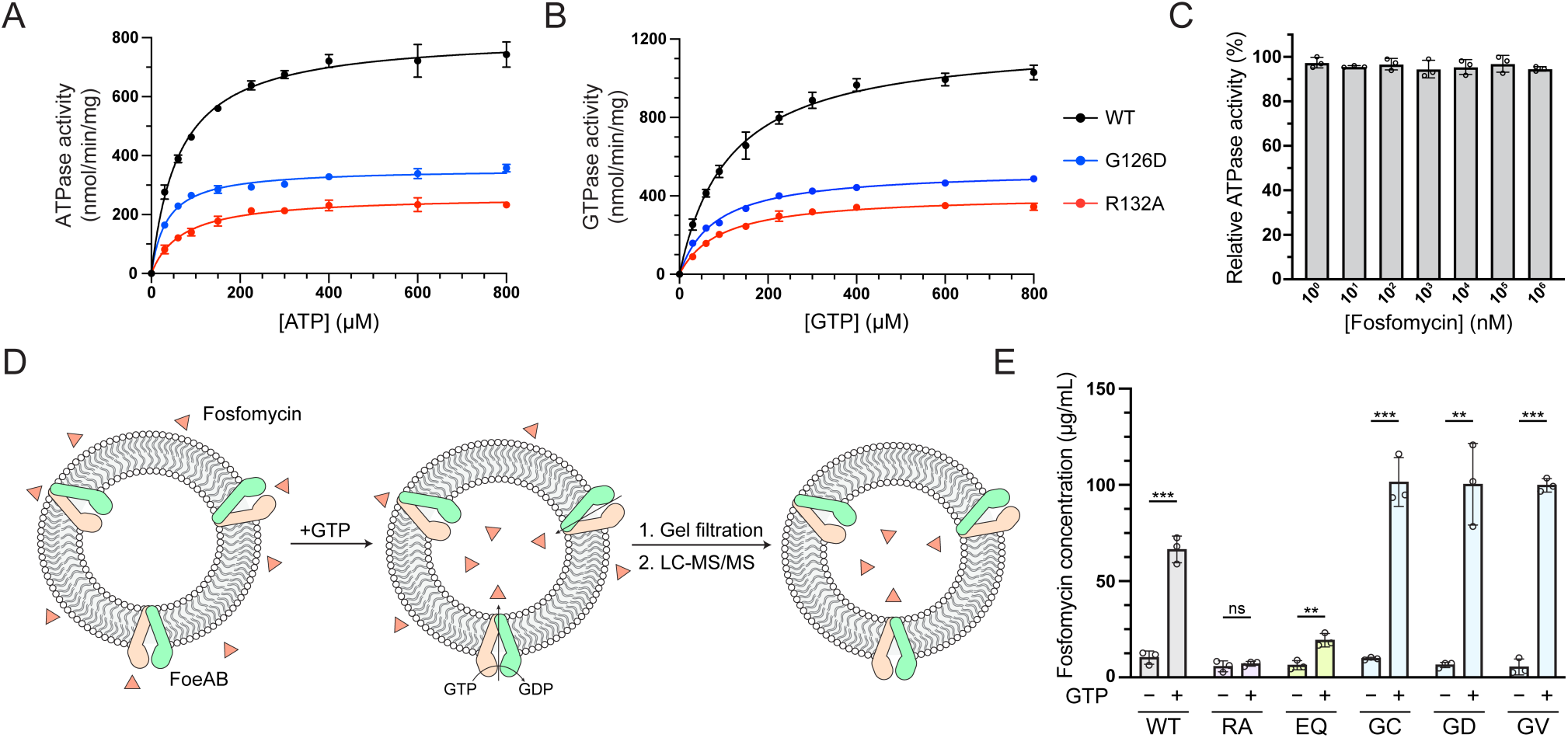
Biochemical reconstitution of FoeAB activity in proteoliposomes. (A) ATPase activity of wild-type FoeAB and the indicated variants. (B) GTPase activity of wild-type FoeAB and the indicated variants. Kinetic parameters are provided in *SI Appendix*, Table S4. (C) Relative ATPase activity of wild-type FoeAB in the presence of increasing concentrations of fosfomycin. Activity was normalized to the basal activity exhibited in the absence of fosfomycin. Data are shown as mean ± SD from triplicates. (D) Schematic of the *in vitro* fosfomycin transport assay using FoeAB-reconstituted proteoliposomes. (E) FoeAB-mediated fosfomycin transport into proteoliposomes in the presence or absence of 5 mM GTP. Fosfomycin transport was quantified by measuring the intraliposomal fosfomycin concentration after 20 min incubation. Data are shown as mean ± SD from triplicates. The effect of GTP addition relative to untreated controls was assessed by two-sided Student’s t-test (\*\**P* < 0.01, \*\*\**P* < 0.001; ns, not significant). Under GTP-treated conditions, fosfomycin transport levels differed significantly between wild-type FoeAB and all variants (*P* < 0.01, one-way ANOVA with Dunnett’s multiple comparison test). RA – R132A^A^; EQ – E504Q^B^; GC – G126C^A^; GD – G126D^A^; GV – G126V^A^.

Next, we sought to evaluate the effect of TMD substitutions on nucleotide hydrolysis, as the fosfomycin resistance phenotype observed in the *St*FoeAB TMD mutants could result from changes in NTPase activity. We selected the hypoactive R132A^A^ and hyperactive G126D^A^ variants for further characterization. The R132A^A^ variant showed a ∼3-fold decrease in both nucleotide hydrolysis rate and catalytic efficiency, indicating that alterations in the positively charged pocket partially impair NTPase activity (Figs. 5A, 5B and *SI Appendix*, Table S4). The decrease was less pronounced for the G126D^A^ mutant; however, because this mutant conferred higher fosfomycin resistance despite lower nucleotide hydrolysis than wild-type *St*FoeAB, changes in turnover rate do not account for the observed phenotype. Likewise, the reduction in NTPase activity alone cannot fully explain the complete loss of fosfomycin resistance in the R132A^A^ mutant, implying that these TMD substitutions primarily affect substrate recognition.

Substrate-dependent modulation of ATPase activity is often used to evaluate substrate specificity of ABC transporters (40). We therefore investigated whether fosfomycin addition affects FoeAB ATPase activity because some prokaryotic ABC exporters are stimulated by substrate addition (38, 41). However, little change in ATPase activity was observed over a wide range of fosfomycin concentrations (Fig. 5C). This result suggests that *St*FoeAB undergoes constitutive nucleotide hydrolysis with little substrate-dependent stimulation.

Having confirmed the NTPase activity of liposome-reconstituted *St*FoeAB, we sought to directly detect fosfomycin transport by *St*FoeAB (Fig. 5D). We prepared a reaction mixture in which fosfomycin was present outside the *St*FoeAB proteoliposomes, and transport was initiated by adding GTP. Because externally added GTP only activates ABC exporters with NBDs oriented towards the exterior of the liposome, this setup enables unidirectional fosfomycin transport into the vesicles. Fosfomycin accumulation was quantified by liquid chromatography tandem mass spectrometry (LC–MS/MS) following removal of external fosfomycin via gel filtration. GTP addition resulted in clear fosfomycin accumulation in proteoliposomes containing wild-type *St*FoeAB, demonstrating that *St*FoeAB transports fosfomycin *in vitro* (Fig. 5E). In contrast, R132A^A^ proteoliposomes showed no appreciable fosfomycin accumulation, further supporting the critical role of this residue in fosfomycin transport. Similarly, fosfomycin transport was severely compromised in the NTPase-impaired E504Q^B^ proteoliposomes, although a modest increase in fosfomycin accumulation was detected, possibly reflecting residual nucleotide hydrolysis activity and/or fosfomycin binding to the transporter. Notably, proteoliposomes containing the G126^A^ variants showed increased fosfomycin accumulation compared with wild-type *St*FoeAB, providing direct evidence that these substitutions enhance fosfomycin transport. Together, these findings establish FoeAB as a fosfomycin exporter and validate the functional importance of residues identified through cellular and structural analyses.

## DISCUSSION

ABC transporters are ubiquitous among living cells including prokaryotes, and their role in drug efflux has long been recognized. In contrast to Gram-negative bacteria, which possess an outer membrane that acts as an additional permeability barrier, Gram-positive bacteria are directly exposed to their surrounding environment, making them more susceptible to xenobiotics. As such, we would expect Gram-positive bacteria to encode multiple ABC exporters that enable the extrusion of harmful compounds. However, only PatAB has previously been described as a multidrug ABC exporter in *S. pneumoniae*, leaving open the question of whether additional pneumococcal ABC exporters contribute to drug efflux. Our work addresses this question through the functional characterization of the newly identified streptococcal ABC exporter FoeAB, which mediates fosfomycin transport—a substrate not previously associated with any prokaryotic efflux pump. Although fosfomycin is not used to treat pneumococcal infections, its use against urinary tract infections caused by enterococcal species with FoeAB homologs raises the possibility that these pumps contribute to reduced fosfomycin susceptibility in clinical settings (22).

The biochemical and structural data presented here enable us to develop a working model of fosfomycin transport that follows an alternating access mechanism. In the basal state, FoeAB adopts an inward-facing conformation that allows fosfomycin to access the putative substrate recognition pocket, where it is coordinated by basic pocket-forming residues. ATP binding to the consensus NBS triggers NBD dimerization and extracellular gate opening, which involves the notable shift of FoeA TM3 that constitutes this pocket. We propose that the dissociation of R136^A^ from R253^B^, which is one of the pocket-forming residues, results in greater conformational flexibility and facilitates substrate dissociation. FoeAB reverts to its inward-facing conformation upon ATP hydrolysis and phosphate release, preparing the transporter for a renewed round of substrate export.

We note that the inward-facing structures of FoeAB in this study may represent a substrate-bound form, as extra density was observed in the positively charged pocket of these structures in the cryo-EM map. This could explain the remarkable similarity in NBD separation among these structures despite differences in their nucleotide binding states. Substrate binding has been shown to facilitate the closure of the inner cleft in some type IV ABC transporters, and it is conceivable that FoeAB forms a wider cavity in the absence of any bound molecules (42, 43). Despite extensive efforts, we were unable to obtain an inward-facing structure lacking this extra density using *E. coli*-expressed FoeAB, which suggests that an alternative expression system may be required to capture a ligand-free inward-facing structure. Furthermore, we anticipate FoeAB to form an occluded state in which both the extracellular and intracellular gates are closed during its conformational cycle, a state that has structurally been captured in other ABC transporters (44, 45). Structural elucidation of these states would provide a more complete picture of FoeAB-mediated substrate export.

While our findings establish FoeAB as a fosfomycin exporter, its ability to transport other substrates remains to be determined. Microbial multidrug ABC exporters typically exhibit high basal ATPase activity that is either unaffected or only slightly activated by the addition of the transported drug(39, 46, 47). Although energetically costly, this feature is thought to confer an advantage by enabling rapid extrusion of toxic compounds without the need for substrate-specific activation (48). FoeAB shares this property, and together with its structural resemblance to known heterodimeric multidrug ABC exporters, current evidence suggests that it may transport anions or phosphonate-/phosphate-containing molecules. However, FoeAB did not confer resistance to fosmidomycin or FR900098, indicating that the presence of a phosphonate group alone is insufficient for substrate recognition. Moreover, phosphorylated metabolites are central to many cellular processes, making their indiscriminate export detrimental to growth. Thus, if FoeAB functions as a multidrug exporter, it likely discriminates substrates by recognizing chemical structures that are less common in cells (*e.g.,* epoxide rings). Future identification of the extra density observed in the positively charged pocket may help clarify the physiological role and substrate specificity of FoeAB. Additional substrates could also be identified through structure-guided *in silico* screening followed by validation using the LC–MS/MS-based transport assay.

This work highlights the potential for greater chemical diversity among the substrates recognized by prokaryotic multidrug ABC exporters, which have traditionally been thought to mediate transport of a similar set of compounds. The majority of efflux pumps in Gram-positive bacteria remain to be characterized, and we anticipate that molecular dissection of their functions will advance our understanding of how these organisms utilize these transporters for their survival and proliferation. Such knowledge could ultimately guide the development of therapeutic approaches aimed at circumventing drug-resistance mechanisms in Gram-positive pathogens.

## EXPERIMENTAL PROCEDURES

### Bacterial culture conditions and strain construction

Culture media were purchased from Becton Dickinson. *E. coli* strains were grown with shaking at 37 °C in lysogeny broth (LB), Terrific Broth (TB), or on LB agar plates with the appropriate additives. *S. pneumoniae* strains were cultured statically in Todd Hewitt broth supplemented with 2% yeast extract (THY) at 37 °C in an atmosphere containing 5% CO_2_. Pre-poured Trypticase Soy Agar with 5% Sheep Blood (TSAII 5%SB) plates with a 5 mL overlay of 1% nutrient broth agar or TSA plates containing 5% defibrinated sheep blood (Japan BioSerum) with appropriate additives were used for growth of pneumococcal cells on solid media. A previously reported protocol was used for *S. pneumoniae* transformation (49). The following concentration of antibiotics were used for selection: carbenicillin, 50 µg/mL; spectinomycin, 100 µg/mL (*E. coli*) or 200 µg/mL (*S. pneumoniae*). Tables summarizing the bacterial strains, plasmids and oligonucleotide primers used in this study as well as protocols for plasmid and strain construction are available in the *SI Appendix*.

### Efflux pump overexpression library screening, viability assay and MIC measurement

*S. pneumoniae* efflux pump overexpression library strains were grown to mid-log phase before adjusting the cell density to OD_600_ = 0.1 in THY. Cells were then diluted 100-fold in phosphate-buffered saline, and 1 µL of the diluted cells were spotted on TSA 5% defibrinated sheep blood plates containing the screening antimicrobial compound. Plates were inspected after overnight incubation for colony formation, and strains that grew in the presence of lethal concentration of antimicrobial compounds were selected for MIC measurement. For viability assay on solid media, the normalized culture was diluted 10-fold four times, and 3 µl dilution sample was spotted on TSA 5% defibrinated sheep blood plates containing the indicated additives. Plates were imaged after overnight incubation.

*S. pneumoniae* liquid MIC was determined by first normalizing mid-log phase cells to OD_600_ = 1. The normalized culture was then diluted 1:500 in THY before adding 100 µL culture to a 96-well plate containing 100 µL THY with the antimicrobial compound of interest. Growth was assessed by measuring absorbance at 600 nm using an Infinite M200 Pro plate reader (Tecan) after overnight incubation. For *E. coli* MIC measurement, cultures grown overnight at 30 °C in LB/carbenicillin were diluted 1:50 in fresh media containing 1 mM IPTG. After 5 h incubation at 30 °C, cells were diluted again in fresh media to OD_600_ = 0.004, and 100 µL of the diluted culture was added to a 96-well plate containing 100 µL LB supplemented with carbenicillin, 50 µg/mL glucose 6-phosphate and 1 mM IPTG. Cell growth was assessed using a plate reader after overnight incubation at 30 °C.

For immunoblot analysis, induced cells were first normalized to OD_600_ = 1, and 30 µL aliquots were mixed with 10 µL 4 x Laemmli sample buffer (Bio-Rad). Proteins were separated by SDS-PAGE using 4-20% polyacrylamide gels, transferred onto nitrocellulose membranes, and blocked with Western BLoT Blocking Buffer (Takara Bio). Membranes were then probed with HRP-conjugated monoclonal anti-DYKDDDDK antibody (1:10000; FUJIFILM Wako Chemicals) or anti-RpoB antibody (1:10000; Abcam), followed by incubation with HRP-conjugated goat anti-rabbit secondary antibody (1:10000; FUJIFILM Wako Chemicals) for anti-RpoB detection. All antibodies were diluted in Tris-buffered saline with 0.1% Tween-20, and chemiluminescent signals were detected using Chemi-Lumi One reagent (Nacalai Tesque).

### Expression and purification of *St*FoeAB

*E. coli* C43(DE3) *ΔacrB* cells harboring the expression plasmid were grown in 1 L TB supplemented with carbenicillin at 37 °C with shaking until OD_600_ ∼0.5. The culture was cooled to 20 °C and protein expression was induced by adding 500 µM IPTG. Cells were harvested 18 h post-induction by centrifugation (4,200 x g, 15 min, 4 °C). All purification steps were performed at 4 °C. Cells were resuspended in 40 mL HBS (20 mM HEPES pH 7.5, 150 mM NaCl) supplemented with 0.25 mg/mL DNaseI and 0.5 mg/mL lysozyme, and the sample was lysed with French press. Cell debris was removed by centrifugation (10,000 x g, 5 min, 4 °C), and the membrane fraction was collected by ultracentrifugation of the supernatant (100,000 x g, 45 min, 4 °C). For preparing protein samples solubilized in DDM, the membrane pellet was resuspended in HBS supplemented with 1% DDM using a glass Dounce tissue grinder (Wheaton). After stirring for 1 h, the solubilized fraction was collected by ultracentrifugation (100,000 x g, 35 min, 4 °C), and the resulting supernatant was applied twice through a 1 mL pre-equilibrated Strep-Tactin 4Flow high-capacity resin (IBA Lifesciences). The resin was washed with 30 mL HBS supplemented with 0.02% DDM (HBS/0.02% DDM) three times, and the protein was eluted in 10 mL HBS/0.02% DDM supplemented with 50 mM biotin. The eluate was further purified by size exclusion chromatography (SEC) with a Superdex 200 Increase 10/300 GL column (Cytiva) equilibrated in HBS/0.02% DDM. Fractions containing the target protein were concentrated by centrifugal filtration. The absorbance at 280 nm was measured using NanoPhotometer NP80 (Implen), and the predicted extinction coefficient was used to calculate protein concentration. FoeAB solubilized in LMNG was purified via the same protocol as above with the following modifications: 0.5% LMNG was used instead of 1% DDM for membrane solubilization, and 0.002% LMNG was used instead of 0.02% DDM in the subsequent steps.

For preparing peptidisc-reconstituted FoeAB, 200 µL FoeAB (5 mg/mL) in HBS/0.02% DDM was mixed with 150 µL peptidisc solution in HBS (10 mg/mL), and the resulting mixture was incubated for 30 min at room temperature. After centrifugation (16,000 x g, 5 min, 4 °C), the supernatant was applied to a Superdex 200 Increase 10/300 GL column equilibrated with HBS, which allowed separation of peptidisc-reconstituted FoeAB from free peptidisc polymers.

### Random mutagenesis screen for isolation of hyperactive *St*FoeAB variants

*St*FoeAB expression plasmid library carrying random point mutations in TM3-6 (R115^A^–V306^A^; V125^B^–L316^B^) was generated by site-saturation mutagenesis of the *St*FoeAB expression plasmid (pATOS167) using primers containing a degenerate NNK codon at the target site. This library was transformed into *E. coli* C43(DE3) *ΔacrB* cells, and transformants were plated on LB agar supplemented with carbenicillin, 50 µg/mL glucose 6-phosphate, 0.1 mM IPTG, and fosfomycin. Colonies that grew on LB plates containing 64 µg/mL or 128 µg/mL fosfomycin were sequenced to identify the mutations introduced in FoeAB. *St*FoeAB expression plasmids harboring these mutations were subsequently reconstructed and transformed into C43(DE3) *ΔacrB* cells for further characterization.

### Cryo-EM sample preparation and data acquisition

For cryo-EM sample preparation, 3-6 mg/mL FoeAB in HBS/0.02% DDM was supplemented with the following additives: nucleotide-free — 4 mM MgCl_2_; WT/ATP — 1 mM ATP and 1 mM MgCl_2_; E504Q/ATP — 1 mM ATP, 1 mM MgCl_2_, 1 mM Na_3_VO_4_ and 10 mM fosfomycin; WT/AMPPNP — 5 mM MgCl_2_, 5 mM AMPPNP and 10 mM fosfomycin; WTpeptidisc/AMPPNP — 1 mM AMPPNP, 1 mM MgCl_2_ and 1 mM fosfomycin. Nucleotides were added immediately before vitrification for all datasets except for the E504Q dataset, where the sample was incubated with ATP at room temperature for 2 min. Samples were applied to a Quantifoil Au R1.2/1.3 200 mesh holey carbon grid or Cu R1.2/1.3 200 mesh holey carbon grid glow-discharged using a JEC-3000FC sputter coater (JEOL) at 20 mA for 20 s or a PIB10 ion bombarder (JEOL) at 10 mA for 1 min. Each grid was blotted for 1-5 s and immediately plunged into liquid ethane using a Vitrobot Mark IV (Thermo Fisher) equilibrated at 4 °C or 18 °C and 100% humidity. All cryo-EM datasets except for the WT/AMPPNP dataset were acquired using SerialEM (v 4.0), yoneoLocr (v 1.0) and JEM-Z300FSC (CRYO ARM 300: JEOL) operated at 300 kV with a K3 direct electron detector (Gatan) in CDS mode (50, 51). The Ω-type in-column energy filter was operated with a slit width of 20 eV for zero-loss imaging. The nominal magnification was 60,000x, corresponding to approximately 0.87 Å per pixel. Defocus varied between −0.5 and −2.0 µm. Each movie was fractionated into 40 frames with a total dose of 80 e^−^/Å^2^. The WT/AMPPNP dataset was acquired using the EPU software and Titan Krios G4 (Thermo Fisher) operated at 300 kV with a K3 direct electron detector (Gatan) in CDS mode. The nominal magnification was 105,000x, corresponding to approximately 0.83 Å per pixel. Defocus varied between −0.8 and −1.8 µm. Each movie was fractionated into 50 frames with a total dose of 50 e^−^/Å^2^.

### Cryo-EM image processing and model building

All datasets except for the WT/AMP-PNP dataset were processed using cryoSPARC (v 4.2.1) (52). A total of 3,820 (nucleotide-free), 13,149 (WT/ATP), 4,711 (E504Q/ATP) and 6,312 (WTpeptidisc/AMPPNP) movies were imported and motion corrected, and the contrast transfer functions (CTFs) were estimated. Micrographs whose CTF max resolutions were beyond 5 Å were selected. To prepare a 2D template, particles were automatically picked from a subset of micrographs using blob picker job and extracted with a box size of 360 pixels binning to 180 pixels. After 2D classification, 2D class averages with the features of secondary structure were selected as templates. A total of 341,371 (nucleotide-free), 3,870,986 (WT/ATP), 602,545 (E504Q/ATP) and 2,749,156 (WTpeptidisc/AMPPNP) particles were automatically picked from all micrographs using the templates and extracted with a box size of 360 pixels binning to 180 pixels. After two rounds of 2D classification, 199,633 (nucleotide-free), 655,928 (WT/ATP), 91,482 (E504Q/ATP) and 357,479 (WTpeptidisc/AMPPNP) particles were selected and extracted with a box size of 360 pixels binning to 300 pixels. The extracted particles were subjected to an ab-initio reconstruction job with three to six classes. For the WTpeptidisc/AMPPNP dataset, heterogeneous refinement was performed using all five classes generated in the ab-initio reconstruction to select the particles classified into a class with higher resolution. A total of 186,084 (nucleotide-free), 430,550 (WT/ATP), 42,885 (E504Q/ATP) and 134,291 (WTpeptidisc/AMPPNP) particles were selected and subjected to non-uniform refinement. After global and local CTF refinement and another round of non-uniform refinement, overall map resolutions (FSC=0.143) reached 2.99 Å (nucleotide-free), 2.84 Å (WT/ATP), 3,16 Å (E504Q/ATP), and 3.08 Å (WTpeptidisc/ATP). For the nucleotide-free dataset, 3D classification without alignment was performed with the mask covering the extramembrane domain, and the classified four classes were subjected to non-uniform refinement separately to reconstruct the final maps at overall resolutions of 3.20 Å (NF-1) 3.26 Å (NF-2), 3.34 Å (NF-3) and 3.25 Å (ADP).

The WT/AMPPNP dataset was processed using RELION-4.0 (53). A total of 6,918 movies were imported and motion corrected, and the CTFs were estimated. Micrographs whose CTF max resolutions were higher than 6 Å were selected. To prepare a 2D template, 3,059,472 particles were automatically picked from micrographs using crYOLO and extracted with a box size of 324 pixels, binning to 108 pixels (54). After 2D classification, 477,058 particles were automatically picked from all 3,059,472 particles, and following three rounds of 3D classification with or without alignment, 245,983 particles were selected and subjected to non-alignment 3D classification. After CTF refinement and Bayesian polishing, overall map resolutions (FSC=0.143) reached 2.91 Å.

The homology model of FoeAB heterodimer was generated using SWISS-MODEL or ColabFold (55, 56). The model was manually modified in Coot (v.0.9.6), refined in PHENIX (v.1.19.2) using real-space refinement, and validated using MolProbity, and this cycle was repeated several times (57–59). Cavity volume was analyzed using the CASTp server using a probe radius of 1.4 Å (60). Molecular graphics were prepared using UCSF Chimera (v.1.16) and PyMOL (v.2.5.8) (61).

### Molecular docking studies

All molecular modeling and docking procedures were carried out using Maestro (v.2021-3; Schrodinger). The three-dimensional structure of the target protein was prepared using the Protein Preparation Wizard, with energy minimization applied only to hydrogen atoms. The three-dimensional structure of fosfomycin was generated using LigPrep with default settings, and the receptor grid for docking was centered on the centroid of R132^A^, R136^A^ and R253^B^. Molecular docking was performed using Glide in standard precision mode with default settings (62).

### Proteoliposome preparation and fosfomycin transport measurement

Proteoliposomes containing *St*FoeAB were prepared by modifying a previously reported LAiR method (63). L-⍺-phosphatidylcholine (10 mg/mL; Sigma Aldrich) in HBS supplemented with 5 mM MgCl_2_ was extruded 11 times through a 200 nm polycarbonate membrane, and 190 µL of the resulting liposome solution was mixed with 10 µL *St*FoeAB (3 mg/mL) in HBS/0.002% LMNG. The mixture was slowly inverted for 30 min at room temperature to facilitate FoeAB reconstitution.

To assess fosfomycin transport activity, 1 mg/mL fosfomycin was added to the 200 µL proteoliposome solution and incubated for 10 min at room temperature. Transport was initiated by adding 5 mM GTP, and the mixture was incubated for 20 min at 37 °C. The resulting mixture was loaded onto a preequilibrated 1 mL Sephadex G-50 resin (Sigma) in Econospin empty size exclusion mini spin column (Epoch Life Science) and centrifuged for 30 sec at 500 x g. The flowthrough was collected and subjected to the same centrifugation step four times to ensure removal of fosfomycin that remained outside of the proteoliposomes. Proteoliposomes were collected by ultracentrifugation (200,000 x g, 30 min, 4 °C) and resuspended in 20 µL ddH_2_O containing the internal standard (0.02 mg/mL fosfomycin-13C_3_; Santa Cruz Biotechnology). The proteoliposome solution was then mixed with 80 µL acetonitrile, and the resulting mixture was vortexed vigorously followed by centrifugation at 21,000 x g for 5 min. The supernatant was passed through an InertSep C18 disposable column (100 mg/1 mL; GL Sciences), and 5 µL of the flowthrough was analyzed by LC-MS/MS on a Waters Xevo TQD mass spectrometer with an electrospray ionization probe operating in negative mode. The sample was separated on a GL Sciences InertSustain Amide HILIC column (3 µm; 2.1 mm x 50 mm) with the following method: flow rate = 0.5 mL/min, 5% solvent A (H_2_O, 10 mM ammonium formate) and 95% solvent B (acetonitrile) for 1 min followed by a linear gradient of solvent B from 95 to 50% over 1 min and 50% solvent B for an additional 1 min. Waters MassLynx Mass Spectrometry software (v. 4.1) was used for analyzing the MS data. Parent ions with *m*/*z* of 137 (fosfomycin) and 140 (fosfomycin-13C_3_) were targeted for MS/MS with a collision energy of 16 eV. Fosfomycin abundance was quantified by measuring the peak size observed for the product ion with *m*/*z* of 79 that was normalized using the product ion signal generated from the internal standard.

### FoeAB NTPase assay

NTPase activity was measured with the EnzChek Phosphate Assay Kit (Invitrogen). All measurements were performed at 25 °C in 100 µL reaction mixture containing 1x reaction buffer (20 mM HEPES pH 7.5, 150 mM NaCl, 5 mM MgCl_2_, 200 µM 2-amino-6-mercapto-7-methylpurine ribonucleoside (MESG) and 0.1 U purine nucleoside phosphorylase) and 0-800 µM NTP. The reaction was initiated by the addition of *St*FoeAB, and NTPase activity was assessed by measuring the increase in absorbance at 360 nm, which corresponds to the amount of released inorganic phosphate, using an Infinite M200 Pro plate reader (Tecan). Kinetic analysis was performed with GraphPad Prism (v.10.4.1) using the Michaelis-Menten model.

## Acknowledgements

We thank Yumiko Takeshita for technical assistance with the experiments. We also thank the cryo-EM facility staff at KEK and Yoichi Sakamaki (University of Tokyo) for assistance with cryo-EM data collection. Funding for this work was provided by the Takeda Science Foundation, Institute for Fermentation (Osaka), Noda Institute for Scientific Research, Nippon Foundation-The University of Osaka Project for Infectious Disease Prevention, JEOL YOKOGUSHI Research Alliance Laboratories of The University of Osaka, MEXT Cooperative Research Program of Network Joint Research Center for Materials and Devices, AMED Research Support Project for Life Science and Drug Discovery (BINDS), AMED iD3 booster grant JP23nk0101653, AMED Basis for Supporting Innovative Drug Discovery and Life Science Research (BINDS) grant JP23ama121003, JP25ama121001 and JP25ama121030, JSPS-NUS Joint Research Program grant JPJSBP120229003 and JSPS KAKENHI grants 21K20759, 23H02631, 23K14518, 23K27322 and 25K02496.

## Competing Interests

The authors declare that they have no conflicts of interest with the contents of this article.

## Data availability

Three-dimensional cryo-EM density maps have been deposited in the Electron Microscopy Data Bank (EMDB) under the accession numbers EMD-66531, EMD-66532, EMD-66533, EMD-66534, EMD-66535, EMD-66536, EMD-66537 and EMD-66953. Atomic coordinates for the atomic models have been deposited in the Protein Data Bank (PDB) under the accession codes 9X46, 9X47, 9X48, 9X49, 9X4A, 9X4B, 9X4C and 9XKA. All other study data are included within the manuscript and/or SI Appendix.

## Author contributions

A.T., J.F., M.T., D.T., K.H., K.F., K.Namba and K.Nishino designed research; A.T., J.F., M.T. and D.T. performed research; A.T., J.F., M.T., D.T., T.M., K.F., K.Namba and K.Nishino analyzed data; and A.T. wrote the paper.

## Notes

### Competing Interest Statement

The authors have declared no competing interest.

