## Supplementary Information for "Structural insights into fosfomycin efflux by a streptococcal ABC transporter"

**Supplementary Methods**

**Plasmid and strain construction**

Efflux pump overexpression plasmids in *S. pneumoniae*: The pPEPZ-$P_{lac}$ vector was first PCR linearized using primers oAT401/oAT402. The linearized vector was ligated to putative efflux pump genes via InFusion Cloning to generate an integrative plasmid that integrates at the pseudogene *spr1750* locus. The following primer pairs were used to PCR amplify the putative efflux pump genes for plasmid construction from *S. pneumoniae* genomic DNA: oAT403/404 (*spr0459-0460*); oAT405/406 (*spr0524-0526*); oAT407/408 (*spr0557-0559*); oAT409/410 (*spr0693-0695*); oAT411/412 (*spr0812-0813*); oAT413/414 (*spr0975-0977*); oAT415/416 (*spr1203-1201*); oAT417/418 (*spr1289-1290*); oAT419/420 (*spr1497-1496*); oAT421/422 (*spr1559-1560*); oAT423/424 (*spr1562-1561*); oAT425/426 (*spr1657-1656*); oAT427/428 (*spr1817-1816*); oAT429/430 (*spr1887-1885*); oAT431/432 (*spr2035-2036*); oAT433/434 (*spr0144*); oAT435/436 (*spr0875*); oAT437/438 (*spr0971*); oAT439/440 (*spr1023*); oAT441/442 (*spr1441*); oAT443/444 (*spr1453*); oAT445/446 (*spr1932*); oAT447/448 (*spr1983*); oAT449/450 (*spr0137*); oAT451/452 (*spr1302*); oAT453/454 (*spr1887-1885*); oAT455/456 (*spr1216-1215*, FoeAB); oAT455/457 (*spr1216*, FoeA); oAT456/458 (*spr1215*, FoeB). To insert *St*FoeAB (*stu0758-0759*) in pPEPZ-$P_{lac}$, primers oAT459/460 was used to amplify the genes from *S. thermophilus* LMG18311 genomic DNA.

*S. pneumoniae* efflux pump overexpression strain construction: *S. pneumoniae* R6 cells were transformed with pPEPZ-$P_{lac}$ plasmid or a DNA cassette containing the putative efflux pump genes. DNA cassettes were prepared by first amplifying pPEPZ-$P_{lac}$ with oAT461/462 and oAT463/464, and these fragments were ligated to efflux genes amplified using primer pairs oAT465/466 (*spr0090*), oAT467/468 (*spr1052*), oAT469/470 (*spr1756*), and oAT471/472 (*spr1908*) by overlapping PCR. Transformants overexpressing the desired efflux pump genes were selected with spectinomycin.

*St*FoeAB expression plasmids: The *stu0758* gene was PCR amplified using primers oAT473/474 and ligated to PCR linearized pETDuet-1 (oAT475/oAT476) via InFusion Cloning. The resulting plasmid was then PCR linearized with oAT477/oAT478 and ligated to PCR amplified *stu0759* (oAT479/oAT480). The plasmid containing both *stu0758* and *stu0759* was then PCR linearized with oAT481/oAT482 and ligated to a TwinStrep tag fragment prepared by PCR amplification with oAT483/oAT484 to generate pATOS167, which expresses *St*FoeA-TwinStrep and *St*FoeB. Primer pairs oAT485/oAT486 ($K22A^A$), oAT487/oAT488 ($R132A^A$), oAT489/oAT490 ($R253A^B$), and oAT491/oAT492 ($E504Q^B$) were used to introduce point mutations in pATOS167. For replacing the TwinStrep tag with a FLAG tag, the expression plasmid was amplified by oAT493/oAT494 and ligated via InFusion Cloning.

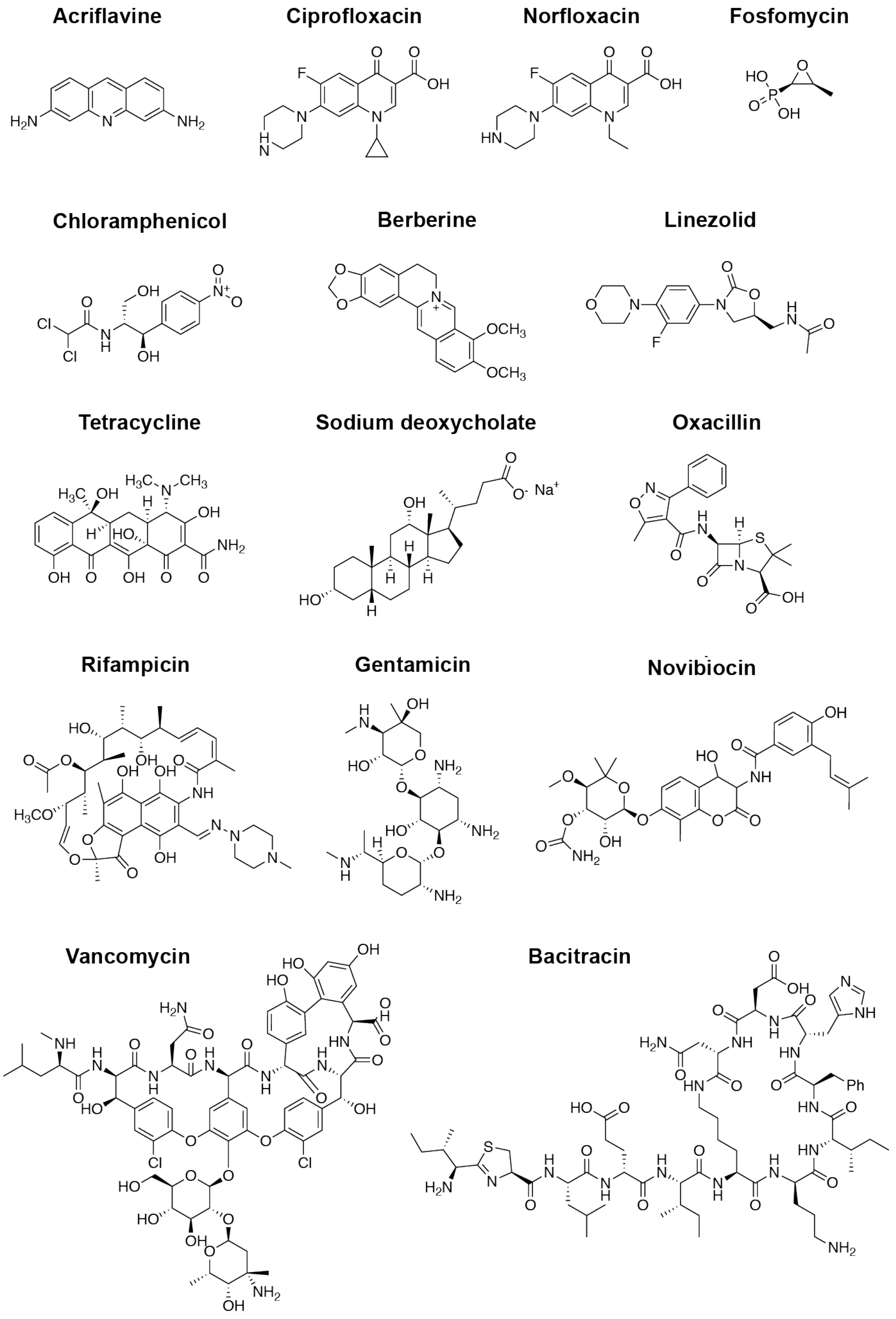


**Figure S1. Chemical structures of antimicrobial compounds screened against the efflux pump overexpression library.** A total of 15 antimicrobial compounds were screened to identify pneumococcal efflux pumps involved in antimicrobial resistance.

A

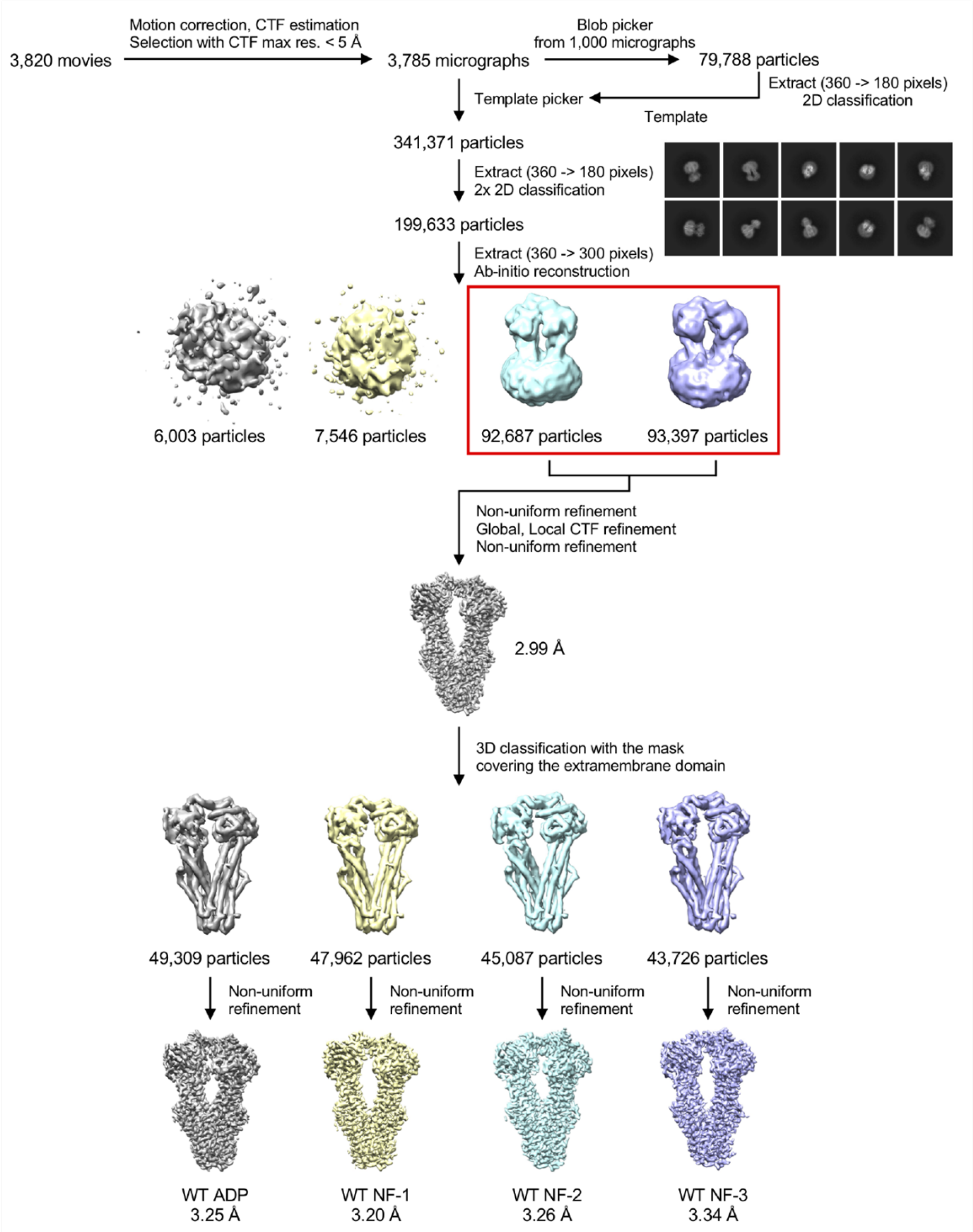
3,820 movies
Motion correction, CTF estimation
Selection with CTF max res. < 5 Å
3,785 micrographs
Blob picker
from 1,000 micrographs
79,788 particles
Extract (360 -> 180 pixels)
2D classification
Template
Template picker
341,371 particles
Extract (360 -> 180 pixels)
2x 2D classification
199,633 particles
Extract (360 -> 300 pixels)
Ab-initio reconstruction
6,003 particles
7,546 particles
92,687 particles
93,397 particles
Non-uniform refinement
Global, Local CTF refinement
Non-uniform refinement
2.99 Å
3D classification with the mask
covering the extramembrane domain
49,309 particles
47,962 particles
45,087 particles
43,726 particles
Non-uniform refinement
Non-uniform refinement
Non-uniform refinement
Non-uniform refinement
WT ADP
3.25 Å
WT NF-1
3.20 Å
WT NF-2
3.26 Å
WT NF-3
3.34 Å

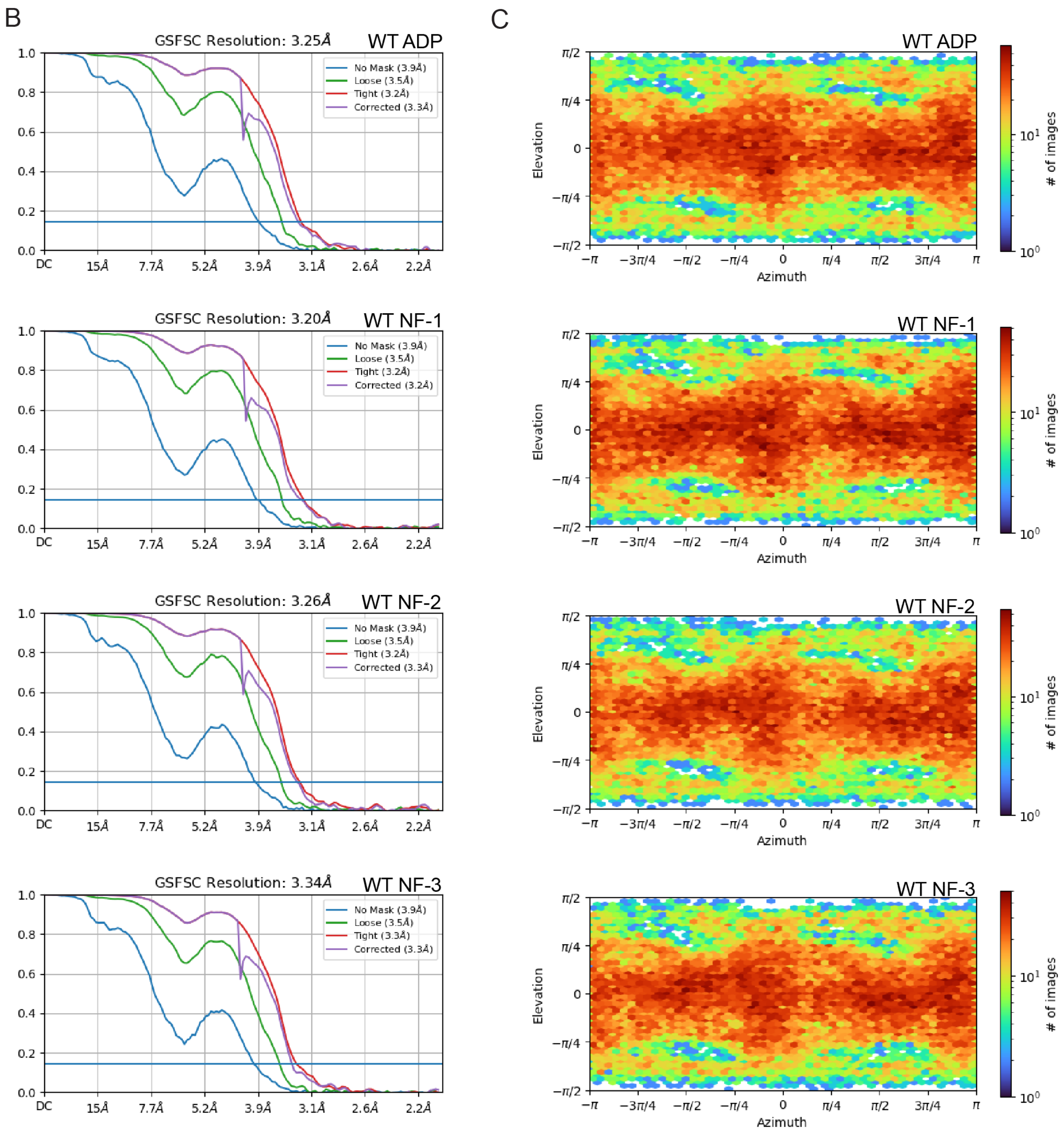


**Figure S2. Cryo-EM data processing of the additive-free dataset.** (A) Cryo-EM data processing workflow using cryoSPARC. (B) Gold-standard Fourier shell correlation (FSC) curves of the EM maps. (C) Angular distribution plots of final particles.

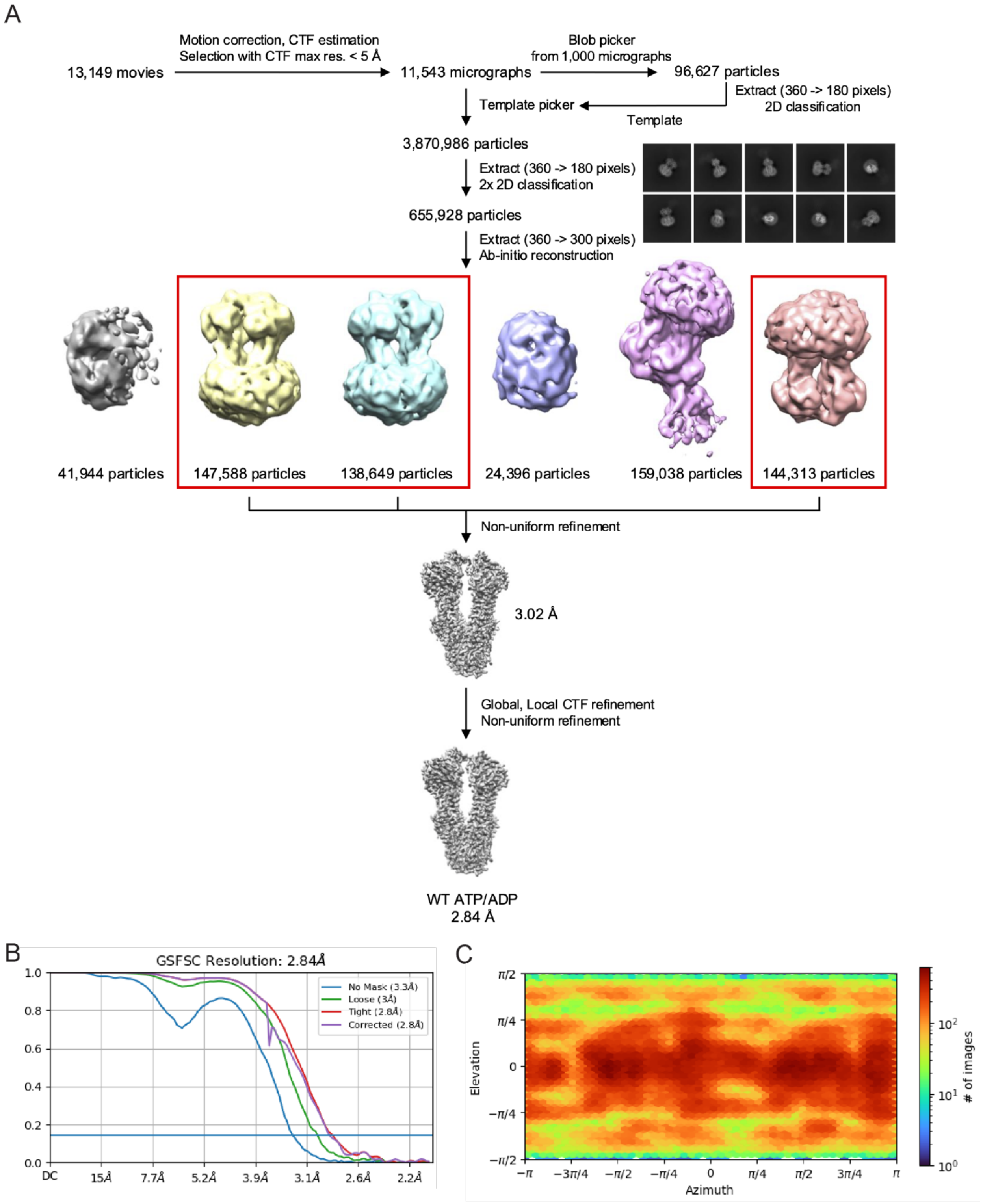


**Figure S3. Cryo-EM data processing of the ATP/ADP-bound FoeAB dataset.** (A) Cryo-EM data processing workflow using cryoSPARC. (B) Gold-standard Fourier shell correlation (FSC) curves of the EM map. (C) Angular distribution plot of final particles.

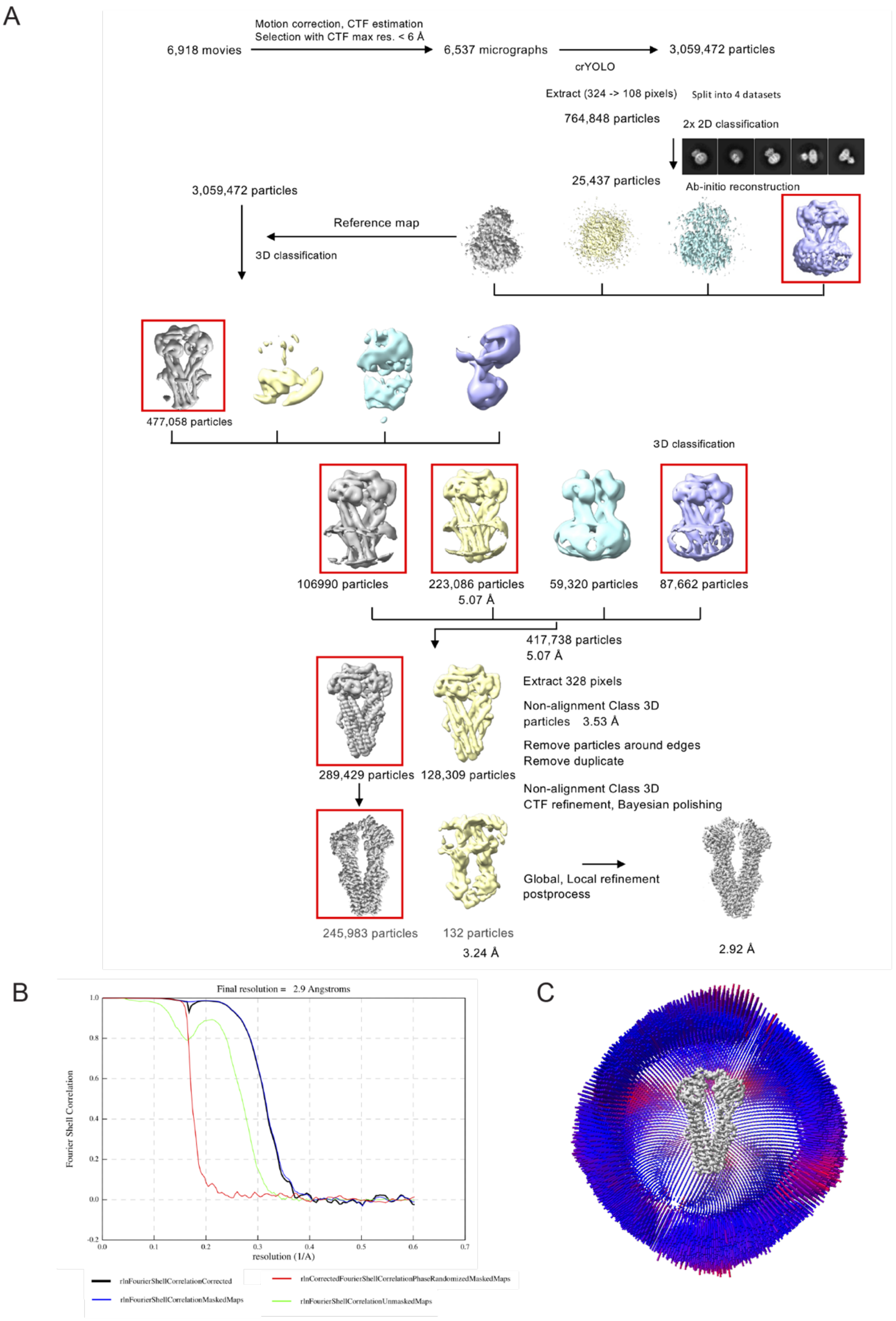


**Figure S4. Cryo-EM data processing of the AMPPNP-bound FoeAB dataset.** (A) Cryo-EM data processing workflow using RELION. (B) Gold-standard Fourier shell correlation (FSC) curves of the EM maps. (C) Angular distribution plots of final particles.

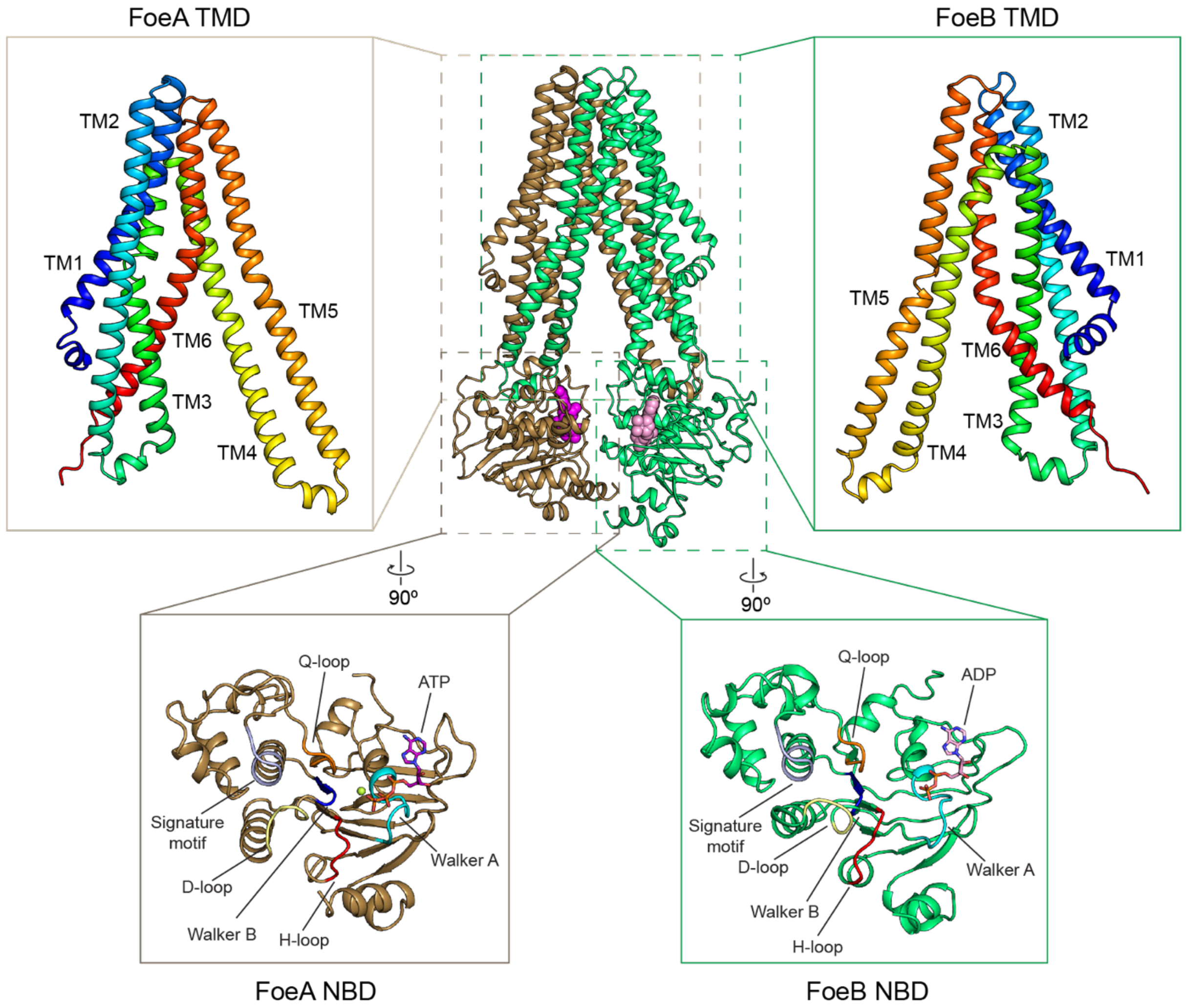


**Figure S5. Domain organization of FoeAB.** Close-up views of the TMDs and NBDs of ATP/ADP-bound FoeAB are shown. Each TMD contains six TM helices, with TM4 and TM5 crossing over and domain-swapping into the opposing protomer. The conserved sequence motifs of the NBDs are individually colored to highlight their spatial organization.

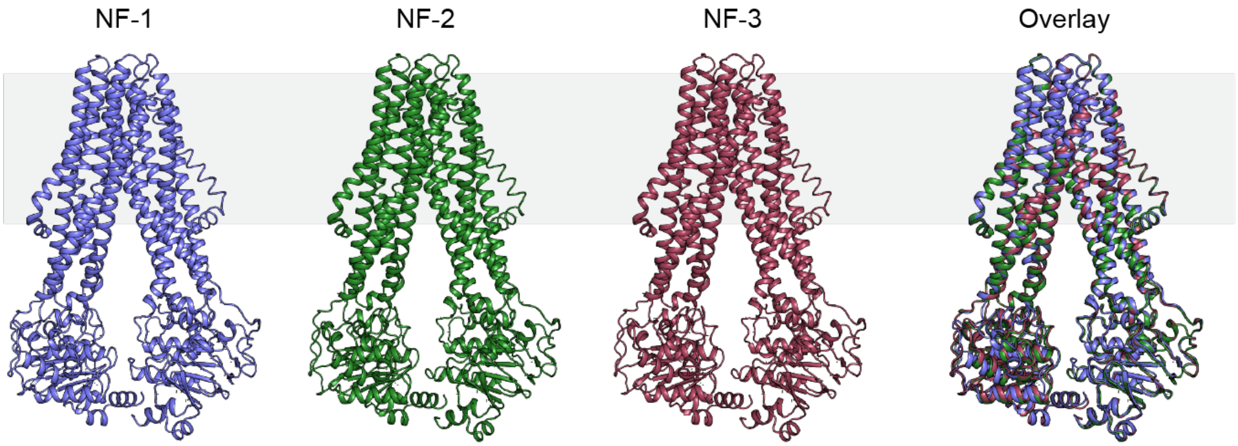


**Figure S6. Inward-facing structures of nucleotide-free (NF) FoeAB.** Nucleotide-free FoeAB structures resolved in this study are shown along with the overlaid structures. Cα root mean square deviation (RMSD) values between these structures are as follows: NF1/NF2 – 0.47 Å; NF1/NF3 – 0.47 Å; NF2/NF3 – 0.17 Å.

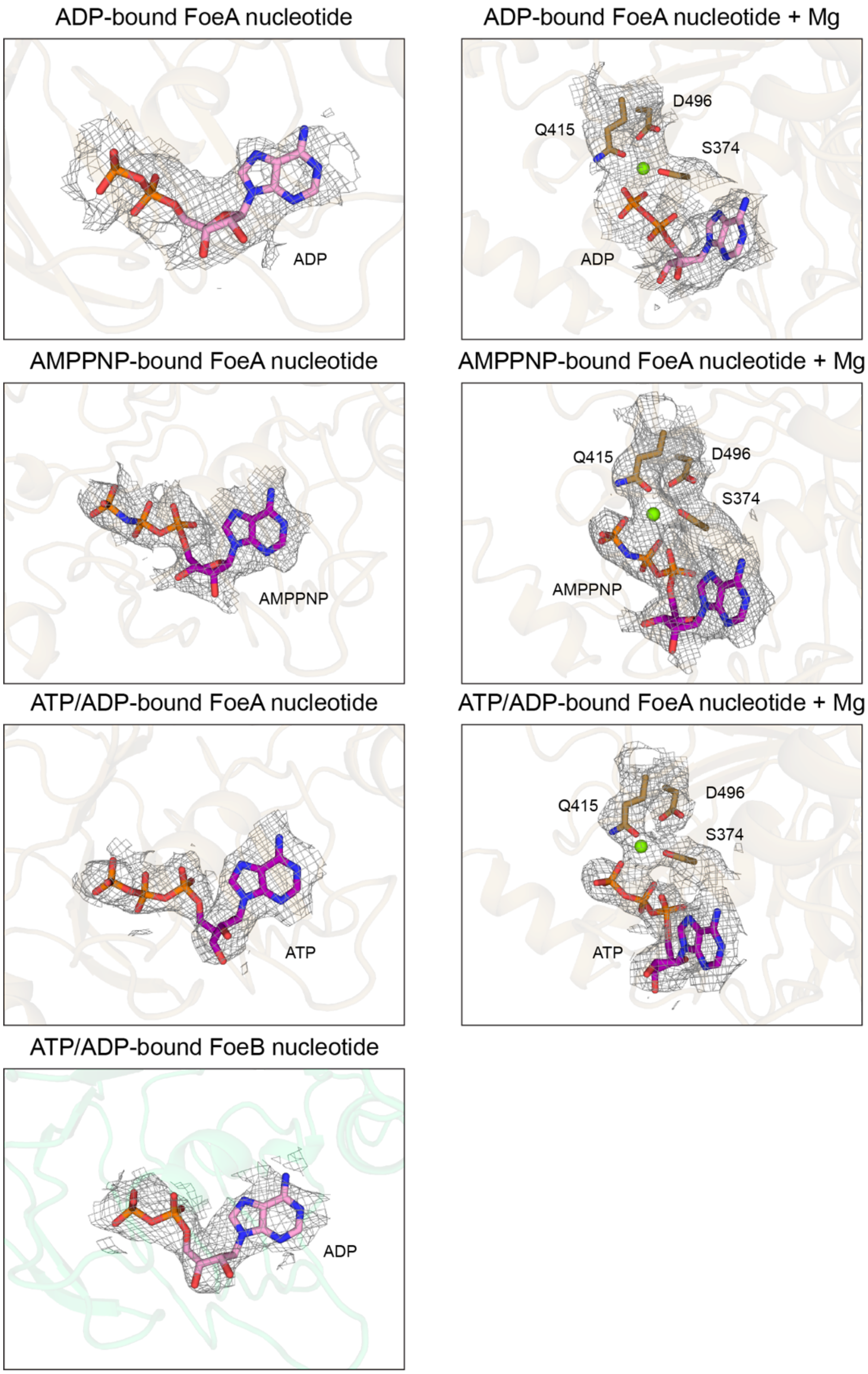


**Figure S7. Cryo-EM densities for bound nucleotides in the inward-facing structures.** Cryo-EM densities corresponding to the nucleotide ligands for ADP-, AMPPNP- and ATP/ADP-bound structures are shown in the left panels. Densities corresponding to the magnesium ion (light green sphere), magnesium coordinating residues, and the bound nucleotide are shown in the right panels. Densities are contoured at 3σ.

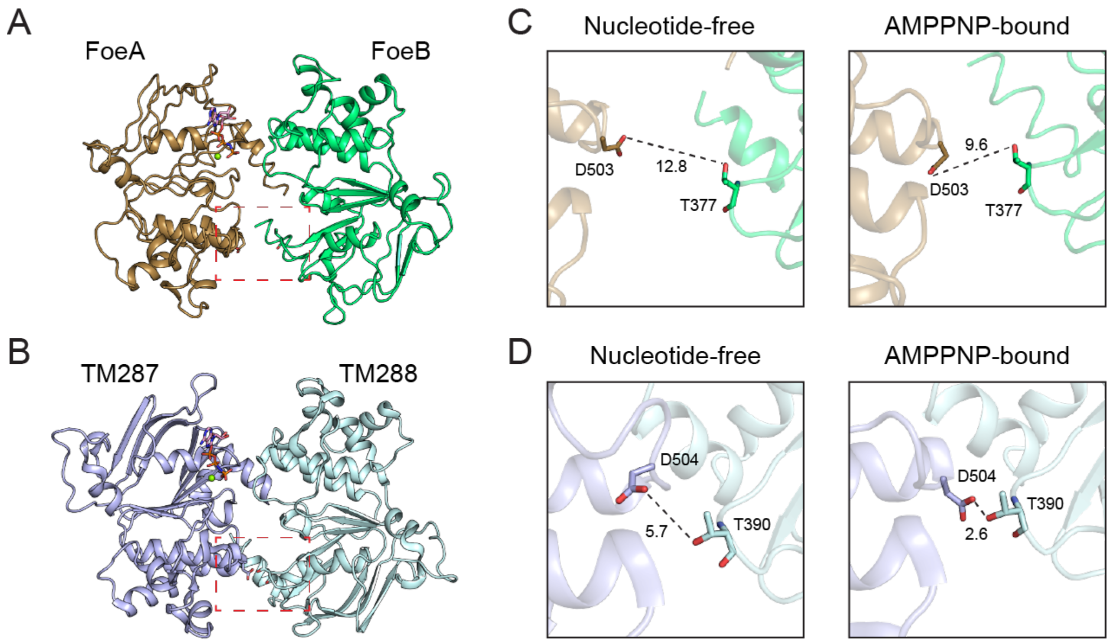


**Figure S8. No direct interaction between the D-loop and Walker A motifs is observed in *St*FoeAB.** (A-B) Top view of the NBDs of FoeAB (A) and TM287/288 (B) AMPPNP-bound structures. The dotted squares indicate the interface between the D-loop and Walker A motifs. (C-D) Close-up views of the interface between the D-loop and consensus Walker A motif in the nucleotide-free and AMPPNP-bound structures of FoeAB (C) and the nucleotide-free (PDB: 4Q4H) and AMPPNP-bound (PDB: 4Q4A) structures of TM287/288. The D-loop aspartate and the Walker A threonine is found to interact only in the TM287/TM288 AMPPNP-bound structure. Numbers denote the distance between these residues in angstroms.

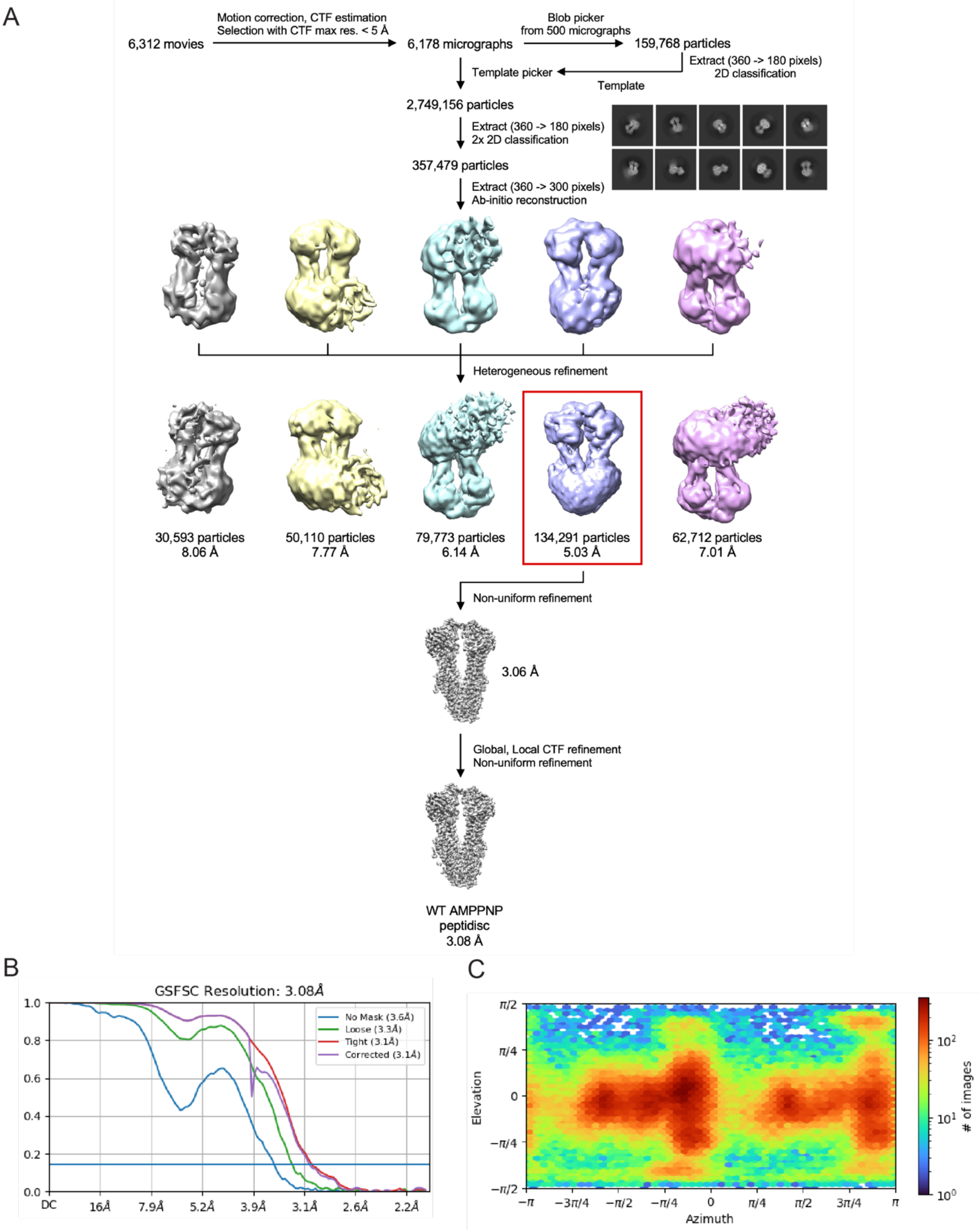


**Figure S9. Cryo-EM data processing of the AMPPNP-bound peptidisc FoeAB dataset.** (A) Cryo-EM data processing workflow using cryoSPARC. (B) Gold-standard Fourier shell correlation (FSC) curves of the EM map. (C) Angular distribution plot of final particles.

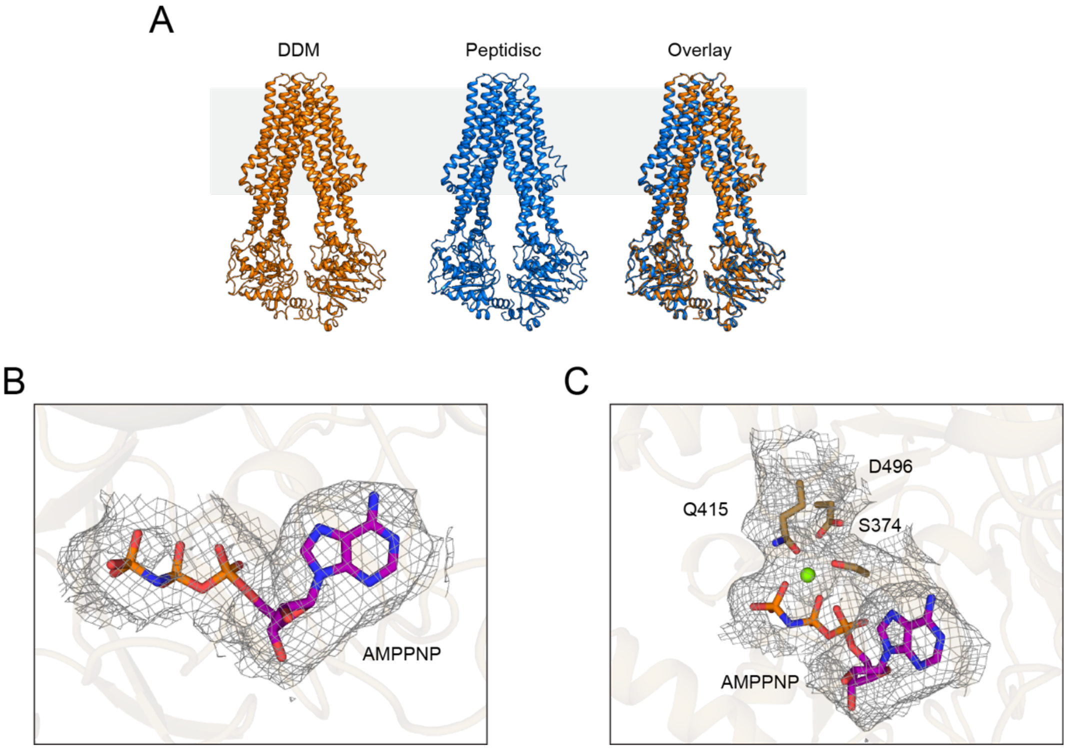


**Figure S10. The use of detergent does not substantially affect FoeAB structure.** (A) AMPPNP-bound FoeAB structures resolved in DDM and peptidisc environments are shown individually along with the overlaid structure. AMPPNP and $Mg^{2+}$ are omitted for clarity. The two structures are nearly identical (RMSD = 0.35 Å) with minimal difference observed in the inter-NBD distance. (B-C) Cryo-EM densities corresponding to AMPPNP and the surrounding residues/magnesium ion in the peptidisc sample. Densities are contoured at 4σ.

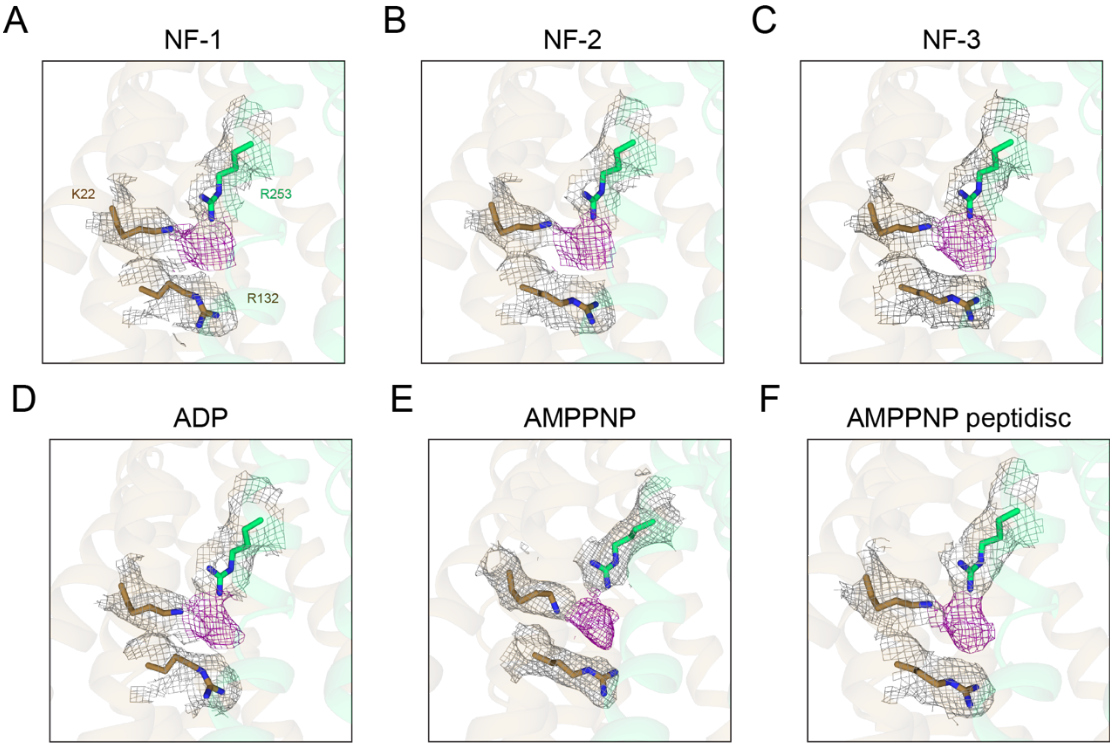


**Figure S11. Extra density for a putative ligand is observed at the positively charged pocket.** Close-up views of the positively charged pocket are shown for nucleotide-free (A-C), ADP-bound (D), AMPPNP-bound (E), and AMPPNP-bound peptidisc (F) structures. Nucleotide-free and ADP-bound structures were obtained in the absence of fosfomycin. Cryo-EM densities corresponding to $K22^A$, $R132^A$ and $R253^B$ are shown in gray, and the extra density attributed to a putative ligand is shown in purple. Densities are contoured at 4σ.

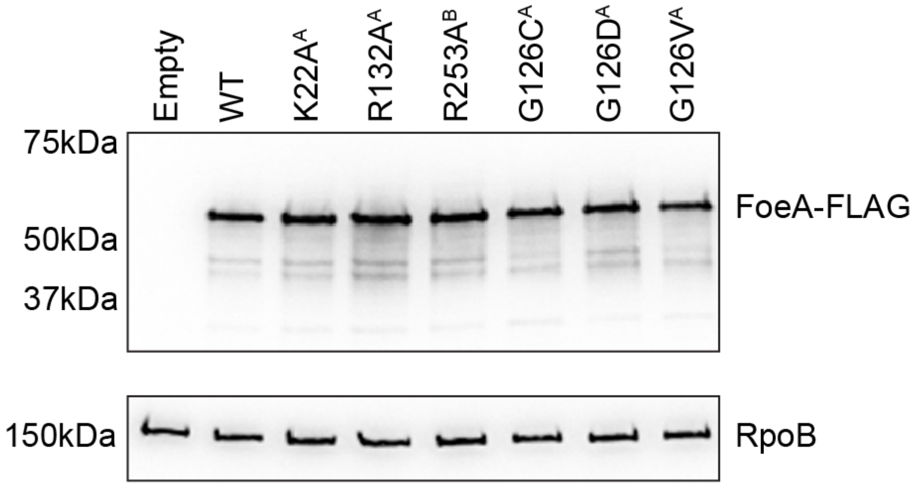


**Figure S12. Western blot analysis of FoeAB expression.** Immunoblot analysis of *St*FoeA-FLAG levels in the *E. coli* strains used in Fig. 3C. *St*FoeA-FLAG was detected using anti-FLAG antibodies, and RpoB was used as a loading control. Representative images from three biological replicates are shown.

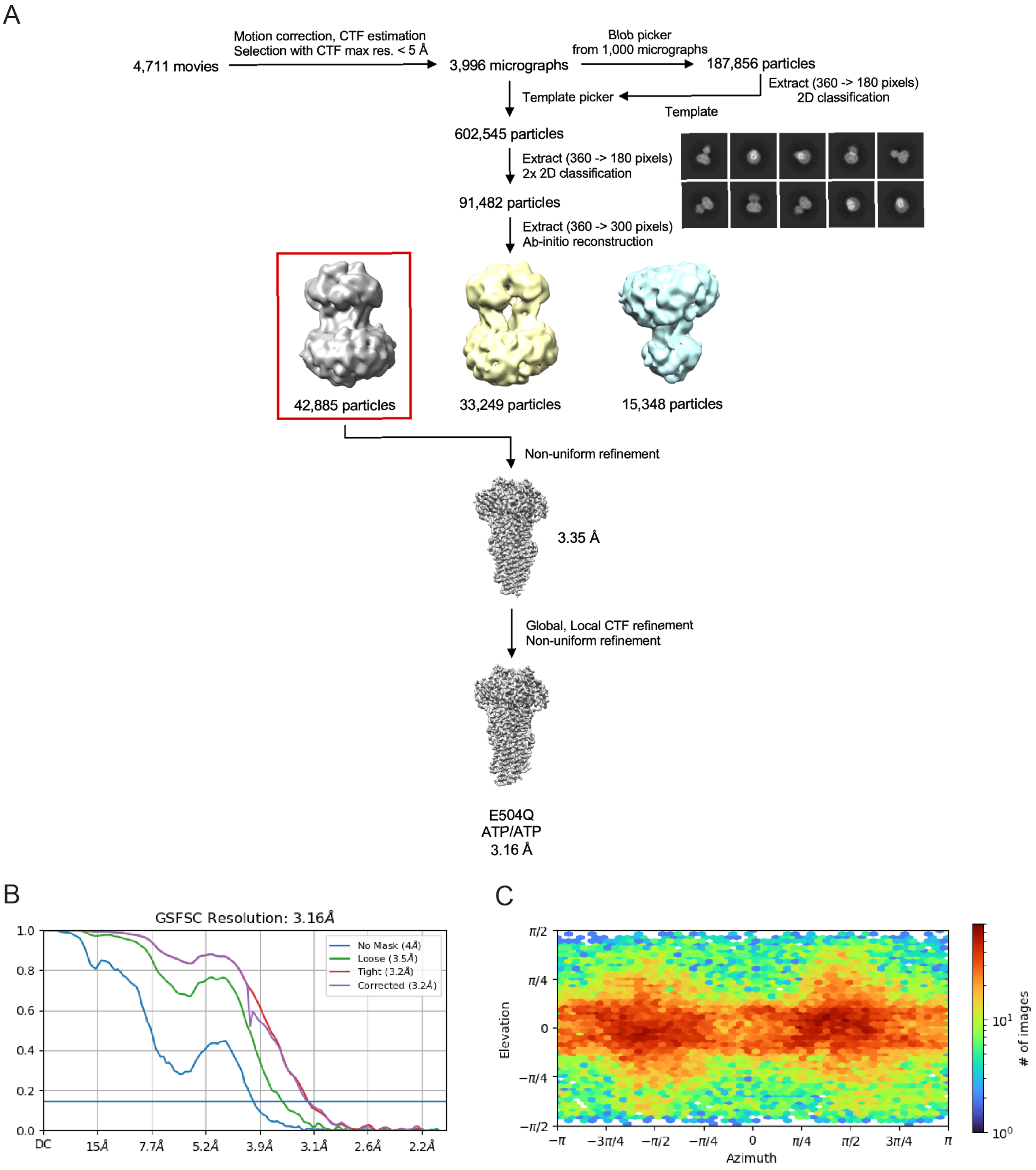


**Figure S13. Cryo-EM data processing of the ATP/ATP-bound FoeAB(E504Q$^{B}$) dataset.** (A) Cryo-EM data processing workflow using cryoSPARC. (B) Gold-standard Fourier shell correlation (FSC) curves of the EM map. (C) Angular distribution plot of final particles.

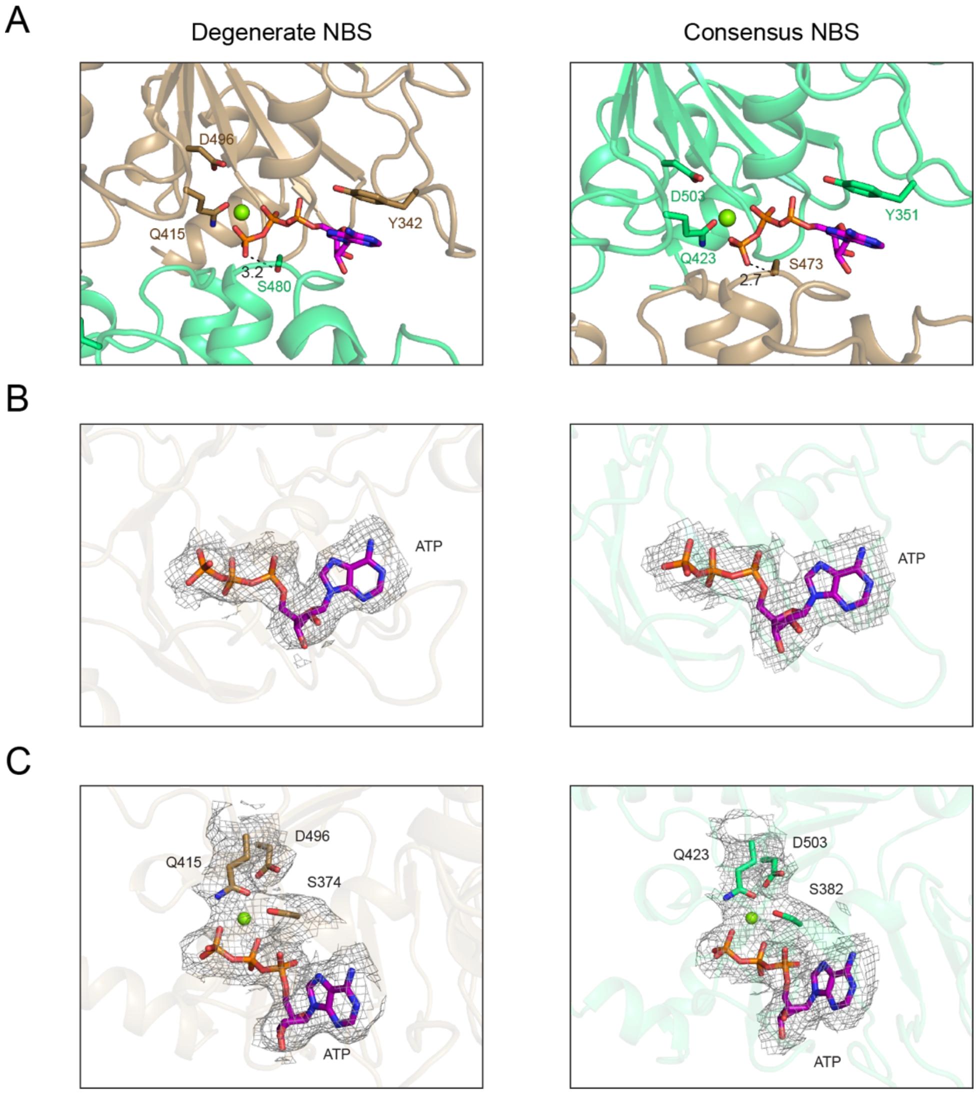


**Figure S14. Close-up views of nucleotide binding sites in the ATP/ATP-bound structure.** (A) The degenerate and consensus NBSs are shown with the two NBDs forming a tight dimer. The distances between the signature motif serine and ATP γ-phosphate are shown in angstroms. (B-C) Cryo-EM densities corresponding to the ATP and the surrounding residues/magnesium ion are shown for the degenerate NBS (left panel) and consensus NBS (right panel). Magnesium ions are depicted as light green spheres. Densities are contoured at 3σ.

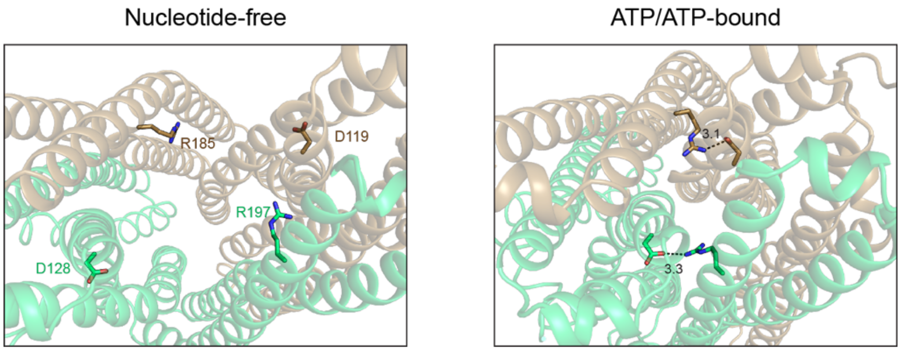


**Figure S15. Intracellular gate closure is stabilized by salt bridges between TM3 and TM4.** A top-down view of the TMD of nucleotide-free and ATP/ATP-bound structures. The outward-facing conformation in the ATP/ATP-bound structure is stabilized by a salt bridge formation between TM3 aspartate and TM4 arginine in both FoeA and FoeB. Numbers denote the distance of salt bridges in angstroms.

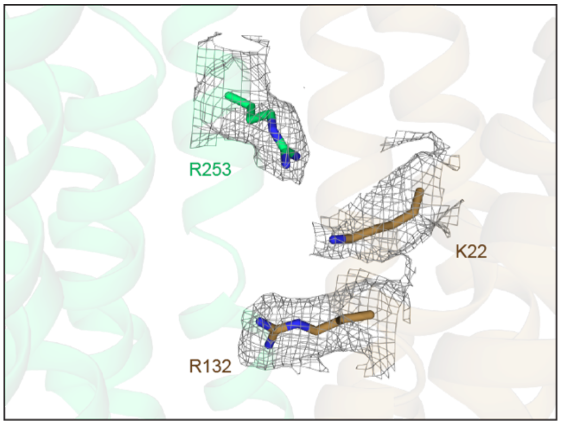


**Figure S16. No extra density is observed in the putative fosfomycin recognition pocket in the ATP/ATP-bound structure.** Density corresponding to the basic residues comprising the pocket is contoured at 3σ.

**Table S1. Putative efflux pump genes investigated in this study.**

| Locus tag | Family | Name | Description | Reference |
|---|---|---|---|---|
| spr0137 | ABC | | Putative transporter | |
| spr0459-0460 | ABC | EcsAB | Putative transporter | |
| spr0524-0526 | ABC | Vex123 | Putative peptide transporter | (1) |
| spr0557-0559 | ABC | | Putative transporter | |
| spr0693-0695 | ABC | | MacAB-like transporter, LL-37 resistance | (2) |
| spr0812-0813 | ABC | BceAB | Bacitracin resistance factor | (3) |
| spr0975-0977 | ABC | | Putative tryptophan/tyrosine transporter | |
| spr1203-1201 | ABC | | Putative transporter | |
| spr1216-1215 | ABC | FoeAB | Fosfomycin transporter | This study |
| spr1289-1290 | ABC | | Putative transporter | |
| spr1497-1496 | ABC | | Putative transporter | |
| spr1559-1560 | ABC | | Putative transporter | |
| spr1562-1561 | ABC | QsrAB | Putative sodium transporter | (4) |
| spr1657-1656 | ABC | | Putative transporter | |
| spr1817-1816 | ABC | | Putative transporter | |
| spr1887-1885 | ABC | PatAB | Multidrug efflux pump | (5, 6) |
| spr2035-2036 | ABC | | Putative transporter | |
| spr0090 | MFS | | Putative cyanate transporter | |
| spr0144 | MFS | VpoC | Putative transporter | |
| spr0875 | MFS | PmrA | Putative multidrug efflux pump | (7, 8) |
| spr0971 | MFS | MefE | Putative macrolide transporter | (9) |
| spr1023 | MFS | | Putative transporter | |
| spr1441 | MFS | OxlT | Putative oxalate/formate antiporter | |
| spr1453 | MFS | | Putative cytidine transporter | (10) |
| spr1932 | MFS | | Putative transporter | |
| spr1983 | MFS | | Putative transporter | |
| spr1052 | MATE | PdrM | Putative multidrug efflux pump | (11) |
| spr1756 | MATE | DinF | Putative transporter | (12) |
| spr1302 | SMR | | Putative transporter | |
| spr1908 | SMR | | Putative transporter | |

| Strain | ACF | BBR | CIP | NOR | TET | BAC |
|---|---|---|---|---|---|---|
| pPEPZ | 16 | 64 | 1 | 4 | 0.25 | 1 |
| PatAB↑ | 32 | 256 | 4 | 16 | 0.5 | - |
| BceAB↑ | - | - | - | - | - | 4 |

**Table S2. Liquid MIC values for *S. pneumoniae* strains overexpressing PatAB or BceAB.** Values are in µg/mL. Strains pPEPZ (empty vector), PatAB↑ and BceAB↑ correspond to AT1104, AT1085 and AT1058, respectively (Table S5). ACF, acriflavine; BBR, berberine chloride; CIP, ciprofloxacin; NOR, norfloxacin; TET, tetracycline; BAC, bacitracin.

### Table S3. Data collection and refinement statistics for *St*FoeAB.

| | WT NF-1 | WT NF-2 | WT NF-3 | WT ADP |
|---|---|---|---|---|
| EMDB ID | EMD-66532 | EMD-66533 | EMD-66534 | EMD-66531 |
| PDB ID | 9X47 | 9X48 | 9X49 | 9X46 |
| **Data collection and processing** | | | | |
| Magnification | 60,000 | | | |
| Voltage (kV) | 300 | | | |
| Electron exposure ($e^{-}/Å^{2}$) | 80 | | | |
| Defocus range (μm) | -0.5 to -2.0 | | | |
| Pixel size (Å) | 1.032 | | | |
| Symmetry imposed | C1 | C1 | C1 | C1 |
| Imported movies | 3,820 | | | |
| Initial particle images | 341,371 | | | |
| Final particle images | 47,962 | 45,087 | 43,726 | 49,309 |
| Map resolution (Å) | 3.20 | 3.26 | 3.34 | 3.25 |
| FSC threshold | 0.143 | 0.143 | 0.143 | 0.143 |
| **Refinement** | | | | |
| Model resolution (Å) | 3.0/3.2/3.4 | 3.1/3.2/3.5 | 3.2/3.3/3.5 | 3.1/3.2/3.4 |
| FSC threshold | 0/0.143/0.5 | 0/0.143/0.5 | 0/0.143/0.5 | 0/0.143/0.5 |
| Model vs Data CC (mask) | 0.85 | 0.85 | 0.85 | 0.86 |
| (volume) | 0.85 | 0.84 | 0.84 | 0.85 |
| Model composition | | | | |
| Chains | 3 | 3 | 3 | 5 |
| Non-hydrogen atoms | 9,041 | 9,041 | 9,041 | 9,069 |
| Protein residues | 1,142 | 1,142 | 1,142 | 1,142 |
| Ligands | - | - | - | 1 (ADP)<br>1 (Mg) |
| R.m.s deviations | | | | |
| Bond lengths (Å) | 0.002 | 0.003 | 0.002 | 0.003 |
| Bond angles (º) | 0.481 | 0.572 | 0.464 | 0.504 |
| Validation | | | | |
| MolProbity score | 1.46 | 1.49 | 1.43 | 1.69 |
| Clashscore | 6.97 | 8.49 | 6.92 | 8.59 |
| Rotamer outliers (%) | 0 | 0 | 0 | 0 |
| Ramachandran plot | | | | |
| Favored (%) | 97.62 | 97.89 | 97.80 | 96.48 |
| Allowed (%) | 2.38 | 2.11 | 2.20 | 3.52 |
| Outliers (%) | 0 | 0 | 0 | 0 |

### Table S3. Data collection and refinement statistics for *St*FoeAB (cont).

| | WT AMPPNP | WT AMPPNP peptidisc | WT ATP/ADP | E504Q ATP/ATP |
|---|---|---|---|---|
| EMDB ID | EMD-66953 | EMD-66537 | EMD-66536 | EMD-66535 |
| PDB ID | 9XKA | 9X4C | 9X4B | 9X4A |
| **Data collection and processing** | | | | |
| Magnification | 105,000 | 60,000 | 60,000 | 60,000 |
| Voltage (kV) | 300 | 300 | 300 | 300 |
| Electron exposure ($e^{-}$/Å$^2$) | 50 | 80 | 80 | 80 |
| Defocus range (μm) | -0.8 to -1.8 | -0.5 to -2.0 | -0.5 to -2.0 | -0.5 to -2.0 |
| Pixel size (Å) | 0.83 | 1.048 | 1.032 | 1.032 |
| Symmetry imposed | C1 | C1 | C1 | C1 |
| Imported movies | 6,918 | 6,312 | 13,149 | 4,711 |
| Initial particle images | 3,059,472 | 2,749,156 | 3,870,986 | 602,545 |
| Final particle images | 245,983 | 134,291 | 430,550 | 42,885 |
| Map resolution (Å) | 2.92 | 3.08 | 2.84 | 3.16 |
| FSC threshold | 0.143 | 0.143 | 0.143 | 0.143 |
| **Refinement** | | | | |
| Model resolution (Å) | 2.4/2.6/2.9 | 3.0/3.0/3.2 | 2.7/2.8/2.9 | 3.0/3.1/3.4 |
| FSC threshold | 0/0.143/0.5 | 0/0.143/0.5 | 0/0.143/0.5 | 0/0.143/0.5 |
| Model vs Data CC (mask) | 0.87 | 0.83 | 0.80 | 0.82 |
| (volume) | 0.84 | 0.79 | 0.75 | 0.83 |
| Model composition | | | | |
| Chains | 5 | 5 | 5 | 4 |
| Non-hydrogen atoms | 9,110 | 9,073 | 9,100 | 9,087 |
| Protein residues | 1,145 | 1,142 | 1,142 | 1,140 |
| Ligands | 1 (AMPPNP)<br>1 (Mg) | 1 (AMPPNP)<br>1 (Mg) | 1 (ATP)<br>1 (ADP)<br>1 (Mg) | 2 (ATP)<br>1 (Mg) |
| R.m.s deviations | | | | |
| Bond lengths (Å) | 0.003 | 0.003 | 0.003 | 0.002 |
| Bond angles (º) | 0.485 | 0.582 | 0.541 | 0.464 |
| Validation | | | | |
| MolProbity score | 1.47 | 1.60 | 1.54 | 1.36 |
| Clashscore | 6.16 | 7.77 | 7.86 | 6.52 |
| Rotamer outliers (%) | 0 | 0 | 0 | 0.10 |
| Ramachandran plot | | | | |
| Favored (%) | 98.07 | 97.01 | 97.45 | 98.24 |
| Allowed (%) | 1.93 | 2.90 | 2.38 | 1.76 |
| Outliers (%) | 0 | 0.09 | 0.18 | 0 |

| Nucleotide | FoeAB | $K_m$ (μM) | $k_{cat}$ ($s^{-1}$) | $k_{cat}/K_m$ ($s^{-1}M^{-1}$) |
|---|---|---|---|---|
| ATP | WT | 63.2 ± 3.7 | 1.80 ± 0.03 | 28481 |
| | G126D[A] | 34.9 ± 2.5 | 0.79 ± 0.01 | 22637 |
| | R132A[A] | 69.0 ± 6.5 | 0.58 ± 0.01 | 8406 |
| GTP | WT | 113.5 ± 6.7 | 2.66 ± 0.05 | 23568 |
| | G126D[A] | 80.5 ± 3.3 | 1.19 ± 0.01 | 14733 |
| | R132A[A] | 91.2 ± 6.6 | 0.90 ± 0.02 | 9877 |

**Table S4. Kinetic properties of *St*FoeAB with respect to ATP and GTP.** All data are mean ± SEM from experiments conducted in triplicate. Kinetic parameters were obtained by fitting the data into the Michaelis-Menten equation.

**Table S5. Bacterial strains used in this study**

| Strain | Description* | Reference |
|---|---|---|
| *E. coli* | | |
| C43(DE3) | Δ*acrB* | (13) |
| *S. pneumoniae* | | |
| R6 | Wild-type | S. Kawabata |
| AT1051 | Δ*spr1750* ::($P_{lac}$- *spr0459-0460*, *spec*); $Spec^R$ | This study |
| AT1052 | Δ*spr1750* ::($P_{lac}$- *spr0524-0526*, *spec*); $Spec^R$ | This study |
| AT1053 | Δ*spr1750* ::($P_{lac}$- *spr0557-0559*, *spec*); $Spec^R$ | This study |
| AT1054 | Δ*spr1750* ::($P_{lac}$- *spr0693-0695*, *spec*); $Spec^R$ | This study |
| AT1058 | Δ*spr1750* ::($P_{lac}$- *spr0812-0813*, *spec*); $Spec^R$ | This study |
| AT1059 | Δ*spr1750* ::($P_{lac}$- *spr0975-0977*, *spec*); $Spec^R$ | This study |
| AT1061 | Δ*spr1750* ::($P_{lac}$- *spr1203-1201*, *spec*); $Spec^R$ | This study |
| AT1063 | Δ*spr1750* ::($P_{lac}$- *spr1289-1290*, *spec*); $Spec^R$ | This study |
| AT1064 | Δ*spr1750* ::($P_{lac}$- *spr1497-1496*, *spec*); $Spec^R$ | This study |
| AT1065 | Δ*spr1750* ::($P_{lac}$- *spr1559-1560*, *spec*); $Spec^R$ | This study |
| AT1066 | Δ*spr1750* ::($P_{lac}$- *spr1562-1561*, *spec*); $Spec^R$ | This study |
| AT1067 | Δ*spr1750* ::($P_{lac}$- *spr1657-1656*, *spec*); $Spec^R$ | This study |
| AT1069 | Δ*spr1750* ::($P_{lac}$- *spr1817-1816*, *spec*); $Spec^R$ | This study |
| AT1070 | Δ*spr1750* ::($P_{lac}$- *spr0144*, *spec*); $Spec^R$ | This study |
| AT1071 | Δ*spr1750* ::($P_{lac}$- *spr0875*, *spec*); $Spec^R$ | This study |
| AT1072 | Δ*spr1750* ::($P_{lac}$- *spr0971*, *spec*); $Spec^R$ | This study |
| AT1073 | Δ*spr1750* ::($P_{lac}$- *spr1023*, *spec*); $Spec^R$ | This study |
| AT1074 | Δ*spr1750* ::($P_{lac}$- *spr1441*, *spec*); $Spec^R$ | This study |
| AT1075 | Δ*spr1750* ::($P_{lac}$- *spr1453*, *spec*); $Spec^R$ | This study |
| AT1076 | Δ*spr1750* ::($P_{lac}$- *spr1932*, *spec*); $Spec^R$ | This study |
| AT1077 | Δ*spr1750* ::($P_{lac}$- *spr1983*, *spec*); $Spec^R$ | This study |
| AT1080 | Δ*spr1750* ::($P_{lac}$- *spr1908*, *spec*); $Spec^R$ | This study |
| AT1081 | Δ*spr1750* ::($P_{lac}$- *spr1052*, *spec*); $Spec^R$ | This study |
| AT1082 | Δ*spr1750* ::($P_{lac}$- *spr1756*, *spec*); $Spec^R$ | This study |
| AT1083 | Δ*spr1750* ::($P_{lac}$- *spr0090*, *spec*); $Spec^R$ | This study |
| AT1084 | Δ*spr1750* ::($P_{lac}$- *spr0137*, *spec*); $Spec^R$ | This study |
| AT1085 | Δ*spr1750* ::($P_{lac}$- *spr1887-1885*, *spec*); $Spec^R$ | This study |
| AT1086 | Δ*spr1750* ::($P_{lac}$- *spr1302*, *spec*); $Spec^R$ | This study |
| AT1087 | Δ*spr1750* ::($P_{lac}$- *spr1216-1215*, *spec*); $Spec^R$ | This study |
| AT1104 | Δ*spr1750* ::($P_{lac}$, *spec*); $Spec^R$ | This study |
| AT1124 | Δ*spr1750* ::($P_{lac}$- *spr2035-2036*, *spec*); $Spec^R$ | This study |

*Abbreviations: $Amp^R$, ampicillin/carbenicillin resistance; $Spec^R$, spectinomycin resistance

### Table S6. Plasmids used in this study

| Plasmid | Description* | Reference |
|---|---|---|
| pPEPZ-$P_{lac}$ | *S. pneumoniae spr1750* integration vector; $Spec^R$ | (14), Addgene |
| pATOS002 | *S. pneumoniae spr1750*::$P_{lac}$-*spr0459-0460* integration vector; $Spec^R$ | This study |
| pATOS003 | *S. pneumoniae spr1750*::$P_{lac}$-*spr0524-0526* integration vector; $Spec^R$ | This study |
| pATOS004 | *S. pneumoniae spr1750*::$P_{lac}$-*spr0557-0559* integration vector; $Spec^R$ | This study |
| pATOS005 | *S. pneumoniae spr1750*::$P_{lac}$-*spr0693-0695* integration vector; $Spec^R$ | This study |
| pATOS009 | *S. pneumoniae spr1750*::$P_{lac}$-*spr0812-0813* integration vector; $Spec^R$ | This study |
| pATOS010 | *S. pneumoniae spr1750*::$P_{lac}$-*spr0975-0977* integration vector; $Spec^R$ | This study |
| pATOS012 | *S. pneumoniae spr1750*::$P_{lac}$-*spr1203-1201* integration vector; $Spec^R$ | This study |
| pATOS015 | *S. pneumoniae spr1750*::$P_{lac}$-*spr1289-1290* integration vector; $Spec^R$ | This study |
| pATOS016 | *S. pneumoniae spr1750*::$P_{lac}$-*spr1497-1496* integration vector; $Spec^R$ | This study |
| pATOS017 | *S. pneumoniae spr1750*::$P_{lac}$-*spr1559-1560* integration vector; $Spec^R$ | This study |
| pATOS018 | *S. pneumoniae spr1750*::$P_{lac}$-*spr1562-1561* integration vector; $Spec^R$ | This study |
| pATOS019 | *S. pneumoniae spr1750*::$P_{lac}$-*spr1657-1656* integration vector; $Spec^R$ | This study |
| pATOS020 | *S. pneumoniae spr1750*::$P_{lac}$-*spr1817-1816* integration vector; $Spec^R$ | This study |
| pATOS023 | *S. pneumoniae spr1750*::$P_{lac}$-*spr2035-2036* integration vector; $Spec^R$ | This study |
| pATOS030 | *S. pneumoniae spr1750*::$P_{lac}$-*spr0144* integration vector; $Spec^R$ | This study |
| pATOS031 | *S. pneumoniae spr1750*::$P_{lac}$-*spr0875* integration vector; $Spec^R$ | This study |
| pATOS032 | *S. pneumoniae spr1750*::$P_{lac}$-*spr0971* integration vector; $Spec^R$ | This study |
| pATOS034 | *S. pneumoniae spr1750*::$P_{lac}$-*spr1023* integration vector; $Spec^R$ | This study |
| pATOS035 | *S. pneumoniae spr1750*::$P_{lac}$-*spr1441* integration vector; $Spec^R$ | This study |
| pATOS036 | *S. pneumoniae spr1750*::$P_{lac}$-*spr1453* integration vector; $Spec^R$ | This study |
| pATOS037 | *S. pneumoniae spr1750*::$P_{lac}$-*spr1932* integration vector; $Spec^R$ | This study |
| pATOS038 | *S. pneumoniae spr1750*::$P_{lac}$-*spr1983* integration vector; $Spec^R$ | This study |
| pATOS050 | *S. pneumoniae spr1750*::$P_{lac}$-*spr0137* integration vector; $Spec^R$ | This study |
| pATOS051 | *S. pneumoniae spr1750*::$P_{lac}$-*spr1302* integration vector; $Spec^R$ | This study |
| pATOS053 | *S. pneumoniae spr1750*::$P_{lac}$-*spr1887-1885* integration vector; $Spec^R$ | This study |
| pATOS096 | *S. pneumoniae spr1750*::$P_{lac}$-*stu0758-0759* integration vector; $Spec^R$ | This study |
| pATOS104 | *S. pneumoniae spr1750*::$P_{lac}$-*spr1216-1215* integration vector; $Spec^R$ | This study |
| pATOS116 | *S. pneumoniae spr1750*::$P_{lac}$-*spr1216* integration vector; $Spec^R$ | This study |
| pATOS117 | *S. pneumoniae spr1750*::$P_{lac}$-*spr1215* integration vector; $Spec^R$ | This study |
| pETDuet-1 | IPTG-inducible protein expression vector containing two multiple cloning sites; $Amp^R$ | Novagen |
| pATOS167 | *St*FoeA-TwinStrep & *St*FoeB expression vector; $Amp^R$ | This study |
| pATOS179 | *St*FoeA-TwinStrep & *St*FoeB(E504Q) expression vector; $Amp^R$ | This study |
| pATOS206 | *St*FoeA(R132A)-TwinStrep & *St*FoeB expression vector; $Amp^R$ | This study |
| pATOS208 | *St*FoeA-TwinStrep & *St*FoeB(R253A) expression vector; $Amp^R$ | This study |
| pATOS211 | *St*FoeA(G126V)-TwinStrep & *St*FoeB expression vector; $Amp^R$ | This study |
| pATOS212 | *St*FoeA(G126D)-TwinStrep & *St*FoeB expression vector; $Amp^R$ | This study |
| pATOS214 | *St*FoeA(G126C)-TwinStrep & *St*FoeB expression vector; $Amp^R$ | This study |
| pATOS219 | *St*FoeA(K22A)-TwinStrep & *St*FoeB expression vector; $Amp^R$ | This study |
| pATOS354 | *St*FoeA-FLAG & *St*FoeB expression vector; $Amp^R$ | This study |

*Abbreviations: $Amp^R$, ampicillin/carbenicillin resistance; $Spec^R$, spectinomycin resistance

**Table S6. Plasmids used in this study (cont.)**

| Plasmid | Description* | Reference |
|---|---|---|
| pATOS356 | *St*FoeA(R132A)-FLAG & *St*FoeB expression vector; $Amp^R$ | This study |
| pATOS357 | *St*FoeA-FLAG & *St*FoeB(R253A) expression vector; $Amp^R$ | This study |
| pATOS358 | *St*FoeA(G126V)-FLAG & *St*FoeB expression vector; $Amp^R$ | This study |
| pATOS359 | *St*FoeA(G126D)-FLAG & *St*FoeB expression vector; $Amp^R$ | This study |
| pATOS360 | *St*FoeA(G126C)-FLAG & *St*FoeB expression vector; $Amp^R$ | This study |
| pATOS361 | *St*FoeA(K22A)-FLAG & *St*FoeB expression vector; $Amp^R$ | This study |

*Abbreviations: $Amp^R$, ampicillin/carbenicillin resistance; $Spec^R$, spectinomycin resistance

**Table S7. Oligonucleotide primers used in this study**

| Primer | Sequence (5′-3′) |
|---|---|
| oAT401 | TATTTTTCCTCCTTATTTATTTAGATCTTAATTGTG |
| oAT402 | GATCCCTCCAGTAACTCGAG |
| oAT403 | TAAGGAGGAAAAATAATGTTAGAAATTAAAAACCTGACAG |
| oAT404 | GTTACTGGAGGGATCTTAGTCCTGCATCTGACGTT |
| oAT405 | TAAGGAGGAAAAATAATGAATCCAATCCAAAGAT |
| oAT406 | GTTACTGGAGGGATCTTATTCACCATCCAGCAAG |
| oAT407 | TAAGGAGGAAAAATAATGGCAATGATAGAAGTGG |
| oAT408 | GTTACTGGAGGGATCTTACGAACCCGCACTTTC |
| oAT409 | TAAGGAGGAAAAATAATGAAGAAAAAGAATGGTAAAGC |
| oAT410 | GTTACTGGAGGGATCTCATTCATAACGAAGGGCTTC |
| oAT411 | TAAGGAGGAAAAATAATGACACTTTTAGATGTAAAACACG |
| oAT412 | GTTACTGGAGGGATCTTACATTTGGACAATCTTACGATAA |
| oAT413 | TAAGGAGGAAAAATAATGTTAAAGGAAATAAAAAGGAGAAAC |
| oAT414 | GTTACTGGAGGGATCTTATTCAAAGAGTTGATAATAATCAGAG |
| oAT415 | TAAGGAGGAAAAATAATGAATTATTTAAAATTTATAAAAAAGACTAAAC |
| oAT416 | GTTACTGGAGGGATCCTATTCCTCCGTAGTTGATTG |
| oAT417 | TAAGGAGGAAAAATAATGAAGATTTACAAAAAACTATTTGC |
| oAT418 | GTTACTGGAGGGATCCTAATAAATAAATTCTTCTGCCATTT |
| oAT419 | TAAGGAGGAAAAATAATGGCTTATATTGAGATGAAACAC |
| oAT420 | GTTACTGGAGGGATCTTACTCTACAGATTTCAGGGC |
| oAT421 | TAAGGAGGAAAAATAATGTCATTACTAGCATTTGAAAATG |
| oAT422 | GTTACTGGAGGGATCTTAAAGCAAATTAACCTTATTTCTC |
| oAT423 | TAAGGAGGAAAAATAATGCTAGAAGTAAGAAGTCTAGAGAAAA |
| oAT424 | GTTACTGGAGGGATCCTATTTATAAGATAAGGCACGTTT |
| oAT425 | TAAGGAGGAAAAATAATGTCCATTATTCAAAAACTTTGG |
| oAT426 | GTTACTGGAGGGATCTCAAAGAGTATAGGCCATGGC |
| oAT427 | TAAGGAGGAAAAATAATGACTGTGGTTAAAGTTGAAAA |
| oAT428 | GTTACTGGAGGGATCTTAAGCATTCTCCTTATGACC |
| oAT429 | TAAGGAGGAAAAATAATGCTGATTCAGAAAATAAAAACCTAC |
| oAT430 | GTTACTGGAGGGATCTTATTCGAAAACAAATTGATTGTGATAG |
| oAT431 | TAAGGAGGAAAAATAATGCTTACAGTATCTGATGTTTC |
| oAT432 | GTTACTGGAGGGATCTTATGCCTTAGATGACTTTCG |
| oAT433 | TAAGGAGGAAAAATAATGAAAGTATTTCTTCAAAATAGAGATTTTA |
| oAT434 | GTTACTGGAGGGATCCTAAATACTTTCACGAATATTCAGAAG |
| oAT435 | TAAGGAGGAAAAATAATGACAGAGATTAACTGGAAGG |
| oAT436 | GTTACTGGAGGGATCCTAGATTTCCTTTACTTTTAATAATGTTC |
| oAT437 | TAAGGAGGAAAAATAATGAAAATAGATAAAAAAAACGAGGC |
| oAT438 | GTTACTGGAGGGATCTCAGCTATTTTTATCATACCTCTTC |
| oAT439 | TAAGGAGGAAAAATAATGAATCGCTATGCAGTGC |
| oAT440 | GTTACTGGAGGGATCTTATATCAATTTTTCAAAATGAGTTCGAAC |

**Table S7. Oligonucleotide primers used in this study (cont.)**

| Primer | Sequence (5′-3′) |
|---|---|
| oAT441 | TAAGGAGGAAAAATAATGAAATCGAATCGTTATATTATTGC |
| oAT442 | GTTACTGGAGGGATCTCATGTAAGCGGTTTTTGACTTTC |
| oAT443 | TAAGGAGGAAAAATAATGAAACAATTTTTAGAACGGGC |
| oAT444 | GTTACTGGAGGGATCCTAAGCTCTCTTCTGAAGTAGGTACAT |
| oAT445 | TAAGGAGGAAAAATAATGAAACTATTGTTTAGAAATCCAGC |
| oAT446 | GTTACTGGAGGGATCTTAGATATCATTTTTGAGATTAAGAATTGTC |
| oAT447 | TAAGGAGGAAAAATAATGTCTAATTCATTTGTCAAGTTG |
| oAT448 | GTTACTGGAGGGATCTCATTTAAAATAATCTCGTCTGATATAAAT |
| oAT449 | TAAGGAGGAAAAATAATGTATACTATTATAAAATCAAATATAAAAAAA |
| oAT450 | GTTACTGGAGGGATCTTAAATAAAATCCTTTGGAAATTGATATATC |
| oAT451 | TAAGGAGGAAAAATAATGTCAAATAGTTTAAAAGGGACTTTAC |
| oAT452 | GTTACTGGAGGGATCTTATTTTTCTAAGAATAAATCTTTCAAAGAAATC |
| oAT453 | TAAGGAGGAAAAATAATGCTGATTCAGAAAATAAAAACCTAC |
| oAT454 | GTTACTGGAGGGATCTTATTCGAAAACAAATTGATTGTGATAG |
| oAT455 | TAAGGAGGAAAAATAATGAAACACCTATTATCTTACTTC |
| oAT456 | GTTACTGGAGGGATCTTATTCAGAACTAAAAGCCGC |
| oAT457 | GTTACTGGAGGGATCCTAGTCCTCCTTTCCATGTT |
| oAT458 | TAAGGAGGAAAAATAATGAAACGACAAACTGTAAAC |
| oAT459 | TAAGGAGGAAAAATAATGAAAGCATTATTAGTTTATTTCAAA |
| oAT460 | GTTACTGGAGGGATCTTAATTATTTTGTGTTTTTTGGGC |
| oAT461 | GCCAATAAATTGCTTCCTTG |
| oAT462 | CATTATTTTTCCTCCTTATTTATTTAGATCTTAATTGTG |
| oAT463 | GATCCCTCCAGTAACTCGAG |
| oAT464 | ATGACACGGATTTTAAGAATAA |
| oAT465 | TAAGGAGGAAAAATAATGAAAAAGCAATCACTCTTTTTTG |
| oAT466 | GTTACTGGAGGGATCTTAAAGGATCTTATCCGCTC |
| oAT467 | TAAGGAGGAAAAATAATGTATAAGACAAAGTGTTTACGAG |
| oAT468 | GTTACTGGAGGGATCTTAGCGTTTGGATTTTGCT |
| oAT469 | TAAGGAGGAAAAATAATGAATAAAAAACGAACAGTGGAC |
| oAT470 | GTTACTGGAGGGATCCTAGGATTGCACTTTGGTTG |
| oAT471 | TAAGGAGGAAAAATAATGAATCAGTATCAGAAAAAGATTG |
| oAT472 | GTTACTGGAGGGATCTTATCTACTTTCTTTCGCTTCTTC |
| oAT473 | AGAAGGAGATATACCATGAAAGCATTATTAGTTTATTTCA |
| oAT474 | TGTTCGACTTAAGCACTAGCCTTCATCAGCGACCA |
| oAT475 | TGCTTAAGTCGAACAGAAAGT |
| oAT476 | GGTATATCTCCTTCTTAAAGTTAAA |
| oAT477 | GGTACCCTCGAGTCTGGT |
| oAT478 | ATGTATATCTCCTTCTTATACTTAA |
| oAT479 | GAAGGAGATATACATATGAAGGCTAGAAATAACAAGTC |
| oAT480 | AGACTCGAGGGTACCTTAATTATTTTGTGTTTTTTGGGC |

**Table S7. Oligonucleotide primers used in this study (cont.)**

| Primer | Sequence (5′-3′) |
|---|---|
| oAT481 | TAGTGCTTAAGTCGAACAGAAAG |
| oAT482 | GCCTTCATCAGCGACCAC |
| oAT443 | GTCGCTGATGAAGGCGGTAGTGCTTGGAGTCATCC |
| oAT484 | TCGACTTAAGCACTACTTCTCAAATTGTGGATGAGACC |
| oAT485 | TGTTTTTGCATTAATCGAAGCTAGTTTCGAACTC |
| oAT486 | ATTAATGCAAAAACAGGTCCCAGGATACTATC |
| oAT487 | GTTTTTAGCACTCTTTTTAAGGGCACCA |
| oAT488 | AAGAGTGCTAAAAACTGATTAATACCTGTTTGA |
| oAT489 | AGCTACAGCTTTTTTTAATAGCTTGCTTTACGCT |
| oAT490 | AAAAAAGCTGTAGCTGGGTTAACCGCT |
| oAT491 | TTTTGGATCAAGCAACATCAAGTATTGATACCCG |
| oAT492 | TTGCTTGATCCAAAATAAGAATTTTAGG |
| oAT493 | ACAAGGACGACGATGACAAGTAGTGCTTAAGTCGAACAGAAAG |
| oAT494 | CATCGTCGTCCTTGTAGTCGCCTTCATCAGCGACCAC |
